# Apical actin filament turnover mediated by cyclase-associated protein is required for organization of non-centrosomal microtubules in epithelium

**DOI:** 10.1101/2025.09.12.675872

**Authors:** Arathi Preeth Babu, Sachin Muralidharan, Konstantin Kogan, Tommi Kotila, Ville Hietakangas, Jaakko Mattila, Minna Poukkula

## Abstract

Epithelial cells rely on the precise coordination of actin and microtubule cytoskeletons to maintain their polarized structure and function. Within polarized epithelial cells, microtubules form longitudinal non-centrosomal arrays essential for organelle positioning and facilitate directional cargo trafficking whereas the apical actin cytoskeleton supports cell shape and tissue architecture. This study investigates the interplay between disrupted apical actin dynamics and microtubule organization *in vivo* within *Drosophila* epithelial tissue. Loss of actin turnover promoting Cyclase-Associated Protein (CAP) results in the excessive accumulation of stable actin structures at the apical cortex. We show that accumulated apical actin sterically excludes microtubules and membrane-bound vesicles. Consequently, non-centrosomal microtubule-dependent processes are impaired, leading to nuclear mispositioning, disrupted apical cargo transport, and defective microvilli formation in CAP mutant cells. Our findings highlight that dense apical actin, due to impaired actin turnover catalyzed by CAP, disrupts the apical organization of non-centrosomal microtubules. This suggests that spatially regulated actin filament turnover is important for microtubule organization and sustaining the polarized functions of epithelial tissues.

## INTRODUCTION

Epithelia are sheets of polarized cells that line tissues and serve as protective barriers against the external environment. These cells exhibit distinct apical and basal surfaces, which enables key epithelial functions such as barrier function, absorption, secretion, and transport (Buckley & St Johnston, 2022). The establishment and maintenance of epithelial cell structure and function depend on the coordinated organization of the actin and microtubule cytoskeletons (Raman et al., 2018). Although, the roles of microtubules in supporting the polarized functions of differentiated epithelial cells are well established (Meiring et al., 2020), the interplay between the actin and microtubules in epithelial cells has been mostly studied in the context of epithelial morphogenesis (Röper, 2020).

The dense apical actin cytoskeletal network is essential for maintaining epithelial cell shape, tissue architecture, and mechanical integrity (Pelaseyed & Bretscher, 2018). The apical, and junctional actomyosin networks, comprising of contractile actin filaments and myosin-2, are key forces driving epithelial morphogenesis (Martin et al., 2009; Roh-Johnson et al., 2012). Additionally, apical actin also supports vesicle trafficking, junctional organization, and cell–cell adhesion. Many epithelial cells feature actin-rich apical structures such as microvilli or stereocilia, that are crucial for tissue functions like nutrient absorption and sensory perception (Sauvanet et al., 2015; Sharkova et al., 2023). The biogenesis of apical microvilli relies on the intracellular apical transport and delivery of membrane vesicles and cargos such as microvilli-specific cadherin PCDH15 / *Drosophila* Cad99C along non-centrosomal microtubules and apical actin network. Defects in this process disrupt microvillar formation (D’Alterio et al., 2005; Khanal et al., 2016; Knowles et al., 2014).

In most epithelia, microtubules are organized into non-centrosomal arrays with their minus-ends anchored at the apical cortex and plus-ends extending toward the basal side (Bartolini & Gundersen, 2006; Meads & Schroer, 1995; Mogensen & Tucker, 1987; Perrin & Matic Vignjevic, 2023; Toya & Takeichi, 2016). This polarized arrangement supports directional trafficking by dynein and kinesin motor proteins, enabling the targeted delivery of vesicles and organelles to apical and basolateral membrane domains, respectively (Kreitzer & Myat, 2018; Reck-Peterson et al., 2018). During epithelial morphogenesis longitudinal microtubule arrays are coordinated with actomyosin contractility (Booth et al., 2014a; Ko et al., 2019a; J.-Y. Lee & Harland, 2007; Singh et al., 2018, 2024; Zhou et al., 2010) The establishment of the non-centrosomal microtubule arrays depends on the apical anchoring of microtubule minus-ends, a process that requires coordinated action of the microtubule-associated proteins CAMSAP3/ *Drosophila* Patronin, and ACF7/MACF1/ *Drosophila* spectraplakin Shot, as well as the apical spectrin cytoskeleton (Khanal et al., 2016; Muroyama et al., 2018; Nashchekin et al., 2016; Ning et al., 2016; Noordstra et al., 2016; Toya et al., 2016). CAMSAP3/Patronin binds, stabilizes, and anchors the minus ends of microtubules at the apical cortex (Goodwin & Vale, 2010; Toya et al., 2016). Shot (ACF7/MACF1) crosslinks microtubules to the actin cytoskeleton, aiding in the assembly and maintenance of non-centrosomal microtubule arrays (Applewhite et al., 2010; S. Lee & Kolodziej, 2002).

It has become increasingly clear that actin and microtubule cytoskeletons interact and are coordinated (Pimm & Henty-Ridilla, 2021). Studies conducted *in vitro* and in cultured cells have revealed that actin structures actively modulate microtubule dynamics and organization over time and space. In particular, the organization and density of filamentous actin not only supports cellular architecture but also has direct consequences for microtubule dynamics and organization, for instance at the leading edge of migrating cells and regulating centrosomal microtubule nucleation during mitosis (Ballestrem et al., 2000; Colin et al., 2018; Farina et al., 2019; Gélin et al., 2023; Gupton et al., 2002; Gupton & Waterman-Storer, 2006; Inoue et al., 2019; Waterman-Storer & Salmon, 1997). This cytoskeletal interplay is especially important for the regulation of cell shape and polarity in migrating cells, and during epithelial morphogenesis (Ko et al., 2019b; Priya, 2022; Ricolo & Araújo, 2020), as well as neuronal and epithelial cells (Dogterom & Koenderink, 2019). However, how regulation of actin dynamics affects structural organization of actin and actin-microtubule coordination, needs to be explored especially in the context of epithelial cell functions *in vivo*.

In this study, we utilize the *Drosophila* follicular epithelium as a model system to investigate the critical role of actin dynamics in the organization and function of non-centrosomal microtubules. Our study demonstrates that actin disassembly protein, Cyclase associated protein-mediated apical actin turnover is required for correct organization of non-centrosomal microtubule arrays in the *Drosophila* follicular epithelium. We show that actin accumulation in CAP deficient cells leads to steric exclusion of non-centrosomal microtubule arrays apically, causing mispositioning of cell nuclei and impairment of apical cargo trafficking crucial for microvilli biogenesis. Together, these findings highlight the importance of actin dynamics for establishment and maintenance of non-centrosomal microtubules and supporting polarized function and structure of epithelial cells.

## RESULTS

### Apical and basal actin filaments are under high turnover in follicular epithelium

To investigate the role of actin dynamics in epithelial cells, we treated *Drosophila* egg chambers (Figure 1 A) with 20 µM Latrunculin A (Lat A). Lat A sequesters actin monomers thus inhibit their polymerization to filamentous actin (Spector et al., 1983). Lat A treatment can be used to monitor the stability of already existing actin filaments. In follicular epithelial cells, Lat A treatment resulted in the disappearance of apical and basal actin, whereas cortical actin remained largely unaffected in both stage 8 and 10 egg chambers (Figure 1B). This suggests that apical and basal actin in follicle cells are under rapid turnover. To investigate the role of apical actin turnover in epithelial cells, we focused on Cyclase-associated protein (CAP aka *Drosophila* Capt), a regulator of actin dynamics. CAP is a well-known actin-binding protein, which promotes rapid actin turnover via its multiple biochemical functions: depolymerization of cofilin-decorated actin filaments, followed by cofilin release and the subsequent recharging of “old” actin monomers by exchanging bound nucleotides from ADP to ATP (Kotila et al., 2018; Lappalainen et al., 2022; Ono, 2013). Notably, loss of CAP leads to an excessive accumulation of filamentous actin specifically at the apical side (Baum & Perrimon, 2001) of *Drosophila* follicular epithelial cells (Figure 1 C; Supl. Figure 1 A). Moreover, Latrunculin A treatment revealed that the accumulated dense actin of CAP mutant cells was very stable as it persisted even after prolonged (2 h) Latrunculin A treatment (Figure 1 D & E).

**Figure 1:**
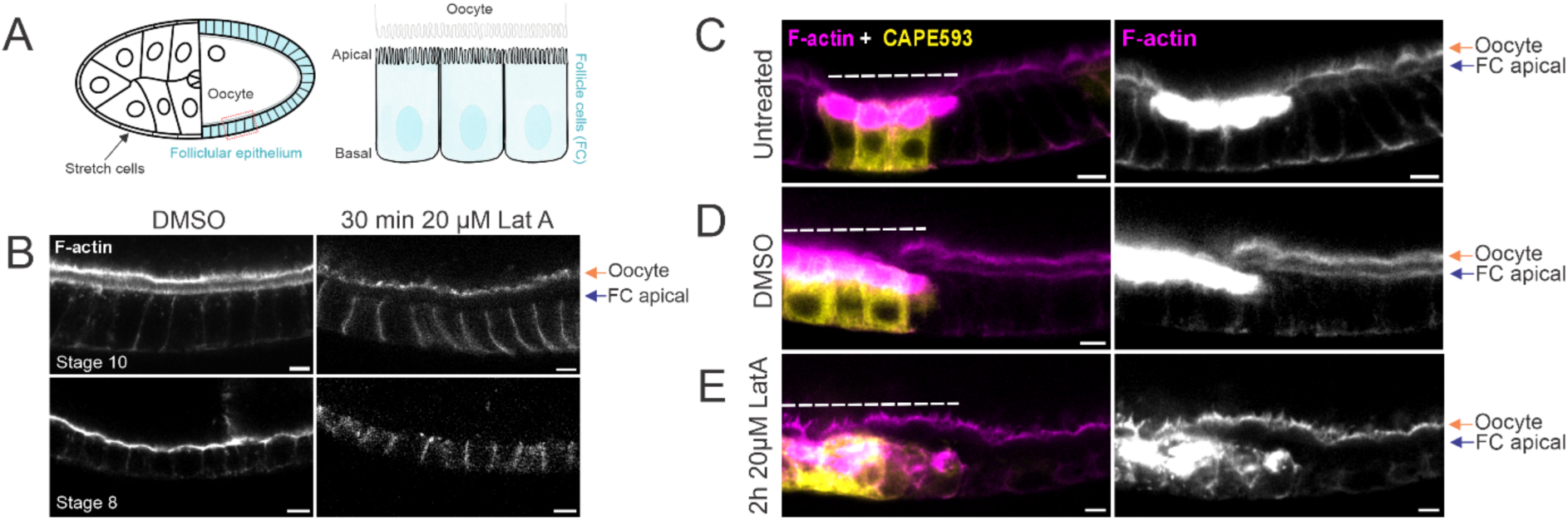
Loss of CAP disrupts fast turnover of apical actin. **(A) Illustration of stage 10 *Drosophila* melanogaster egg chamber** highlighting the apicobasal polarization of the follicular epithelium. The black arrow points towards the squamous cells called stretch cells. **(B) Latrunculin A treatment reveals filamentous actin structures that are under fast turnover.** Treatment of egg chambers with DMSO (control) and 20 µM Latrunculin A for 30-minutes results in a decrease in filamentous actin structures under fast turnover. In stage 10 follicular epithelium, filamentous actin (stained with Phalloidin) is decreased in apical microvilli, apical actin cortex, and basal stress fibres, while junctional actin and the oocyte actin cortex appear to be less affected. In stage 8 follicular epithelium, Latrunculin A induces depletion of both apical actin and basal actin in the follicle cells, while junctional actin remains unaffected. The orange arrow highlights the oocyte cortex, and the blue arrow highlights apical surface of the follicle cells. Scale bar = 10 µm. **(C) Loss of CAP results in apical actin accumulation.** Mosaic follicular epithelium with CAP mutant cells (GFP, pseudo-colored yellow, dotted line) exhibiting significant apical actin accumulation (Phalloidin, pseudo-colored magenta). Scale bar = 10 µm. **(D-E) Even prolonged Latrunculin A treatment failed to fully disassemble dense accumulated actin of CAP mutant cells.** Mosaic follicular epithelium with CAP mutant cells marked with mCD8-GFP (GFP positive, pseudo-colored yellow, dashed line), stained for filamentous actin (stained with phalloidin, pseudo-colored magenta) treated with DMSO (control) **(D)** and 20 µM Latrunculin A **(E)** for 2 hours. Upon treatment with Latrunculin A, apical actin accumulation persisted in CAP mutant cells whereas a decrease in apical F-actin was observed in the neighbouring control cells. Scale bar = 10 µm.

Despite highly localized accumulated actin, CAP protein itself displayed mostly diffuse cytoplasmic localization (Supl. Figure 1 A). In other systems (Bertling et al., 2004; Ono, 2013), CAP targets specifically those actin filament populations that are under rapid turnover. We investigated if specific actin filament configurations would be enriched in CAP mutant cells. Indeed, accumulated apical actin in CAP mutant cells displayed increased levels of proteins promoting rapid actin filament disassembly, such as Aip1 and Twinfilin (Supl. Figure 1 B, D), suggesting compensatory mechanisms for managing excess of filamentous actin. While the actin filament elongator Ena was enriched in these actin structures (Supl. Figure 1 E; Baum & Perrimon, 2001), profilin, which supplies monomeric actin for elongation, was expressed at higher levels but was not localized at the site of actin accumulation (Supl. Figure 1 C). This indicates that an excess of actin filament assembly is unlikely to occur at the apical side of CAP mutant cells. Additionally, α-actinin and tropomyosin were enriched at the site of actin accumulation (Supl. Figure 1 F-G), whereas active non-muscle myosin II was either partially or completely excluded from apical actin networks in a stage-dependent manner, suggesting that the accumulated actin is unlikely to form contractile actomyosin filaments (Supl. Figure 1 H, H’). In conclusion, the dense and stabilized actin in CAP mutant cells is composed of a complex mixture of filamentous actin and various actin-binding proteins but appears to lack components providing contractile properties.

### Nucleotide exchange activity of CAP is required for maintaining proper apical actin dynamics

CAP is a multidomain protein, with each domain exhibiting distinct biochemical activities that contribute to actin turnover (Figure 2 A). The N-terminal Helical Folded Domain (HDF) enhances depolymerization of cofilin-decorated actin filaments by dissociating cofilin-actin monomers (Bertling et al., 2004; Chaudhry et al., 2013; Jansen et al., 2014; Kotila et al., 2019; Quintero-Monzon et al., 2009; Shekhar et al., 2019). The Polyproline (PP) domain binds profilin (Bertling et al., 2007). The Wiskott-Aldrich Syndrome Protein Homology 2 (WH2) domain interacts with both ADP-actin and ATP-actin monomers (Bertling et al., 2007; Freeman et al., 1995; Lila & Drubin, 1997; Mattila et al., 2004). The CARP domain binds ADP-actin monomers and facilitates nucleotide exchange from ADP to ATP, thereby recharging actin monomers for polymerization (Bertling et al., 2004; Kotila et al., 2018; Moriyama & Yahara, 2002).

**Figure 2.**
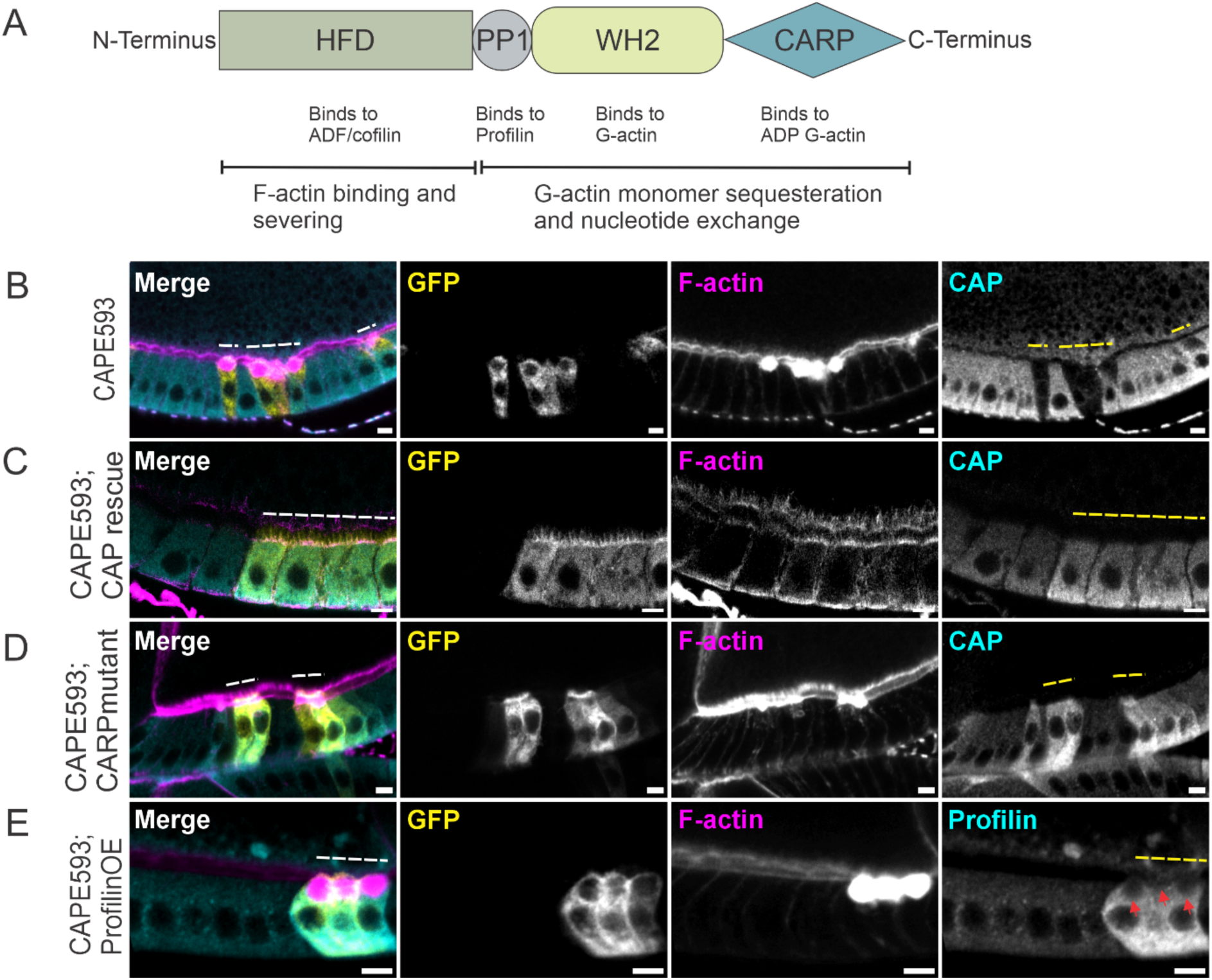
Nucleotide exchange activity of CAP is critical for apical actin turnover. **(A) Schematic diagram of protein domain organization of CAP. (B) CAP mutant cells shows apical actin accumulation**. Mosaic follicular epithelium with CAP mutant cells (GFP, pseudo-colored yellow, dashed line) exhibiting significant apical actin accumulation (Phalloidin, pseudo-colored magenta) alongside a loss of CAP expression (immunostaining, pseudo-colored cyan). Scale bar = 10 µm. **(C) CAP rescue restores apical actin accumulation.** Mosaic follicular epithelium with CAP mutant cells rescued with wildtype CAP (GFP positive, pseudo-colored yellow, dashed line) demonstrating restored apical actin accumulation (Phalloidin, pseudo-colored magenta) and expression of the CAP rescue construct (immunostaining, pseudo-colored cyan). Scale bar = 10 µm. **(D) CARP domain mutant fails to rescue apical actin accumulation.** Mosaic follicular epithelium with CAP mutant cells rescued with a CARP domain mutant of CAP (GFP positive, pseudo-colored yellow, dashed line) showing failure to restore apical actin accumulation (Phalloidin, pseudo-colored magenta). Expression of the CAP in CARP-domain rescue construct is depicted (immunostaining, pseudo-colored cyan). Scale bar = 10 µm. **(E) Profilin overexpression fails to rescue apical actin accumulation.** Mosaic follicular epithelium with CAP mutant cells overexpressing profilin (GFP positive, pseudo-colored yellow, dashed line) demonstrating failure to restore apical actin accumulation (Phalloidin, pseudo-colored magenta). The expression of the profilin construct is shown (immunostaining, pseudo-colored cyan). The profilin expression is increased at the basal region of the cells. Red arrows denote the actin accumulation site lacking profilin localisation. Scale bar = 10 µm.

The existing biochemical, structural, and cell biological studies have contributed to a comprehensive understanding of how CAP’s biochemical activities are orchestrated to promote actin turnover. Therefore, to understand how CAP mediates apical actin turnover *in vivo*, we generated mutant constructs targeting critical amino acids in the HFD, PP, WH2, and CARP domains (Figure 2 A, Table 1) and tested the ability of these constructs to rescue the CAP mutant phenotype. Our results show that CAP constructs with mutations in HFD, PP, and WH2 domains rescued the actin accumulation phenotype in stages 8-10 of oogenesis (Supl Figure 2 B-E). However, the CARP domain mutant did not rescue actin accumulation in the CAP mutant cells even in stages 9-10 (Figure 2 B-D). This result shows that the nucleotide exchange activity of CAP is most critical for efficient apical actin turnover. Our observation is consistent with the fact that nucleotide exchange is the most conserved function of CAP across eukaryotes (Ono, 2013) and its loss leads to significant actin accumulation in yeast (Kotila et al., 2018). Another established nucleotide exchange factor for actin is profilin, and in budding yeast the loss of CAP can be rescued by over-expression of profilin (Vojtek et al., 1991). However, in CAP mutant follicle cells over-expression of profilin did not rescue apical actin accumulation (Figure 2 E). The failure of profilin overexpression to rescue the CAP mutant phenotype suggests that merely increasing nucleotide exchange is not sufficient to restore proper actin dynamics in these cells, indicating a more complex regulatory mechanism at play. Surprisingly, a rescue construct with mutations in the N-terminal domain (responsible for promoting the depolymerization of cofilin-decorated actin filaments) effectively rescued the CAP mutant phenotype. Efficient actin depolymerization typically requires multimerization mediated by the CAP N-terminus (Chaudhry et al., 2013). However, *Drosophila* CAP lacks the critical N-terminal sequence for multimerization (Benlali et al., 2000), which may explain this finding. In conclusion, our findings highlight the critical role of CAP’s nucleotide exchange activity in regulating actin dynamics particularly in the apical region.

**Table 1:** CAP domain mutations.

| Domains | Mutation | Reference | Function |
| --- | --- | --- | --- |
| HFD<br>(90)<br>Interface1 | F122A, Y123A | (Kotila et al., 2019; Quintero-Monzon et al., 2009)(Kotila et al., 2019; Quintero-Monzon et al., 2009)(Kotila et al., 2019; Quintero-Monzon et al., 2009) | Cofilin-actin interaction<br>Pointed end depolymerization |
| PP | <sup>179</sup>PPPPPPP<sup>190</sup><br>→ <sup>179</sup>AAAAAAA<sup>190</sup> |  | Profilin binding |
| WH2<br>(101) | K225A K226A | (Mattila et al., 2004) | ATP-actin binding |
| CARP<br>(m6) | K315A N317A D322A | (Kotila et al., 2018) | ADP-actin binding,<br>nucleotide exchange |

### CAP mutant cells display abnormal distribution of membrane-bound organelles

In our experiments we used MARCM system with mCD8-GFP as a clonal marker to outline the mutant cells in chimeric tissues (T. Lee & Luo, 2001). The membrane-bound mCD8-GFP labels cell membranes and apical microvilli (Blanc et al., 1988; Burr et al., 2014). In follicle cells, mCD8-GFP marks also the entire cytoplasm, possibly highlighting the enrichment of endomembrane compartments in these cells. Interestingly, in CAP mutant cells mCD8-GFP-labeled compartments were lost at the regions of actin accumulation (Figure 3 A). The phenotype was rescued by reintroducing wild-type CAP (Figure 3 B). This result suggests that accumulated actin causes significant impairment in the normal distribution of endomembrane components in CAP mutant cells. Moreover, we found that the site of actin accumulation was devoid of membrane-bound organelles, indicated by the absence of specific markers for the endoplasmic reticulum (ER marker calnexin Cnx99A) and Golgi (trans-Golgi-localizing Golgin 84) (Figure 3 C, D). Transmission electron microscopy (TEM) imaging of CAP mutant cells confirmed that the site of actin accumulation lacked all membrane-bound cytoplasmic components (Figure 3 E i, ii). Additionally, CAP mutant cells displayed ectopic electron-dense vesicular structures both beneath and surrounding the site of actin accumulation (Figure 3 E iii, iv). Notably, also Gutzeit, (1986) described appearance of electron-dense structures in follicle cells upon disruption of microtubules and suggested that these vesicular structures contain vitelline membrane components destined for apical secretion. Highly active apical secretion by follicle cells during stages 9-10 is required for depositing extracellular components and vitelline membrane proteins into the extracellular space between oocyte and follicle cells. This space progressively widens to form the perivitelline space, a membrane component of mature eggshell. Interestingly, the width of the perivitelline space, apically to CAP mutant cells was decreased compared to the neighboring cells (Figure 3 E), further providing evidence that apical secretion of vitelline membrane components in CAP mutant cells was impaired. In conclusion, apically accumulated dense actin in CAP mutant cells excludes endomembrane cytoplasmic components and vesicles destined for apical secretion and formation of perivitelline space.

**Figure 3.**
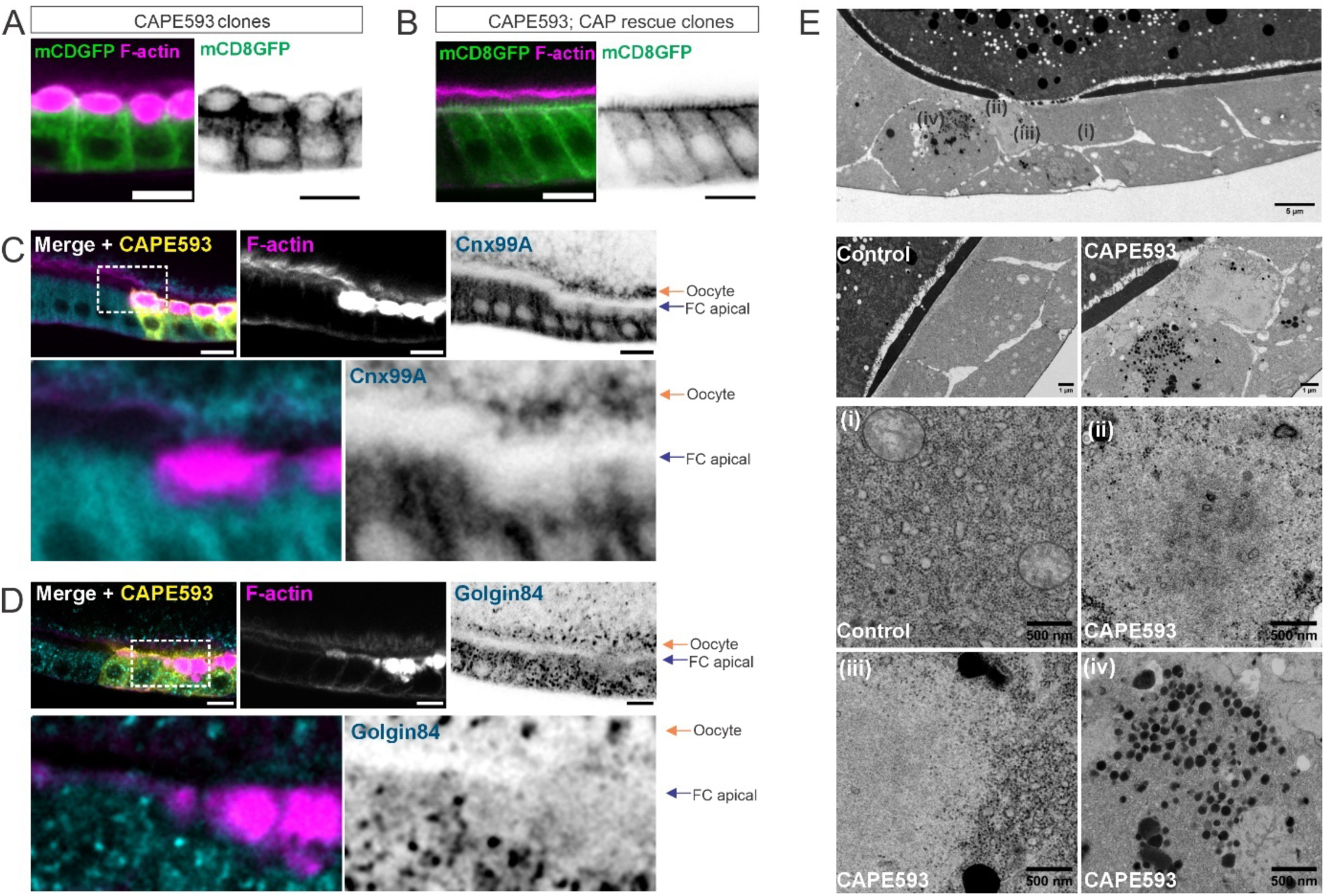
Membrane-bound organelles are excluded from site of actin accumulation. **(A) Absence of membrane-bound mCD8-GFP at site of actin accumulation in CAP mutant cells.** Mosaic follicular epithelium with CAP mutant cells marked with mCD8-GFP (GFP positive, green) and filamentous actin (stained with phalloidin, pseudo-colored magenta). In CAP mutant cells, the membrane bound mCD8-GFP clonal marker (green) is absent in the region of actin accumulation (magenta). Scale bar = 10 µm. **(B) mCD8-GFP localization is restored upon CAP rescue. Mosaic follicular epithelium expressing wildtype CAP in CAP mutant cells.** CAP rescue (mCD8-GFP positive cells) restores the actin accumulation phenotype (F-actin, magenta) and re-establishes mCD8GFP (green) localization. Scale bar = 10 µm. **(C) The site of actin accumulation lacks ER markers.** Mosaic follicular epithelium with CAP mutant cells (GFP positive, pseudo-colored yellow) stained for F-actin (phalloidin, pseudo-colored magenta) and antibody against ER marker Calnexin99A (Cnx99A, pseudocoloured, cyan). Zoomed-in inset shows the apical region of the cell with a lack of ER markers at the site of F-actin accumulation. The orange arrow highlights the Cnx99A localisation on the oocyte cortex, and the blue arrow highlights the localisation of Cnx99A on the apical surface of the follicle cells (FC). Scale bar = 10 µm. **(D) The Golgi marker is lacking at the site of actin accumulation.** Mosaic follicular epithelium with CAP mutant cells (GFP positive, pseudo-colored yellow) stained for F-actin (phalloidin, pseudo-colored magenta) and antibody against Golgi marker Golgin84 (pseudocoloured, cyan). Zoomed-in inset shows the apical region of the cell with F-actin accumulation, lacking the Golgi marker Golgin84. The orange arrow highlights the Golgin84 localisation on the oocyte cortex, and the blue arrow highlights the localisation of Golgin84 on the apical surface of the follicle cells (FC). Scale bar = 10 µm. **(E) Transmission electron microscopy (TEM) of CAP mutant cells.** Transmission electron microscopy (TEM) image of follicular epithelium with CAP mutant clones and control cells. The site of actin accumulation appears less dense than other regions of the cell and neighboring control cells and lacks organelles such as mitochondria, ER, and ribosomes. The numbered regions correspond to specified magnified insets. **(i)** Magnified region near the apical domain of the control cell, showing the presence of membrane-bound organelles like mitochondria. **(ii)** Magnified image of the region near the apical domain in CAP mutant cells, showing the lack of cytoplasmic components. **(iii)** Magnified image of the boundary of the site of actin accumulation, highlighting the difference in cytoplasmic components inside and outside of the accumulated actin. **(iv)** Magnified image showing the presence of electron-dense vesicular structures near the site of actin accumulation in CAP mutant cells.

### Defects in CAP-mediated apical actin turnover affect formation of apical microvilli

The formation of the perivitelline space begins during mid-oogenesis (stages 8–9), when follicle cells develop dense apical microvilli that interdigitate with microvilli projecting from the oocyte. Formation of apical microvilli has been shown to be required for proper expansion of the perivitelline space (Khanal et al., 2016). Therefore, we measured the length of apical microvilli and found them to be significantly shorter in CAP mutant cells compared to controls or mutant cells rescued with wild type CAP (Figure 4 A, B). A well-known component of follicle cell microvilli and a crucial regulator of their biogenesis is protocadherin Cad99C, a *Drosophila* homolog of PCDH15 (Protocadherin-15) (D’Alterio et al., 2005; Schlichting et al., 2005). In wild-type cells, Cad99C was localized at the apical microvilli (Figure 4 C). However, in CAP mutant cells, the localization of Cad99C at the apical surface was reduced, with most of the protein retaining beneath the site of actin accumulation (Figure 4 C, E-F). This result shows that the dense apical actin networks in CAP mutant cells prevent trafficking of Cad99C-containing vesicles to the apical surface. Cad99C is an established cargo transported by Dynein and Rab11 in follicle cells (Khanal et al., 2016). Therefore, we investigated whether microtubule-based transport is disrupted in CAP mutant cells. Transport of apical cargoes are mediated by the microtubule minus-end-directed motor protein Dynein (Harris & Peifer, 2005; Horne-Badovinac & Bilder, 2008; Reck-Peterson et al., 2018; Wilkie & Davis, 2001), which carries Rab11-positive recycling endosomes toward the apical cortex, where microtubule minus-ends are anchored. Rab11, a member of the RAB GTPase family, is critical for polarized trafficking of various cargoes, including membrane proteins and junctional components (Casanova et al., 1999; Jing & Prekeris, 2009; Roeth et al., 2009). We first examined the localization of Dynein and Rab11-positive endosomes in the CAP mutant follicle cells. In control cells, both Dynein and Rab11 were enriched at the apical surface. However, in CAP mutant cells, Dynein and Rab11 localized beneath the accumulated apical actin (Figure 4 F–H’, I–K’). The localization of Rab11 beneath accumulated actin suggest that trafficking of Dynein and Rab11-mediated cargoes to the apical cortex is impaired by the excess apical actin in CAP mutant follicle cells. It should be noted that apical trafficking of Cad99C is also mediated by Myosin V motor protein along the apical actin network (Khanal et al., 2016), but our current experiments do not address whether its functions are affected in CAP mutant cells. Taken together, these results demonstrate that CAP-mediated apical actin turnover is needed for trafficking of apical cargoes and for microvilli biogenesis.

**Figure 4.**
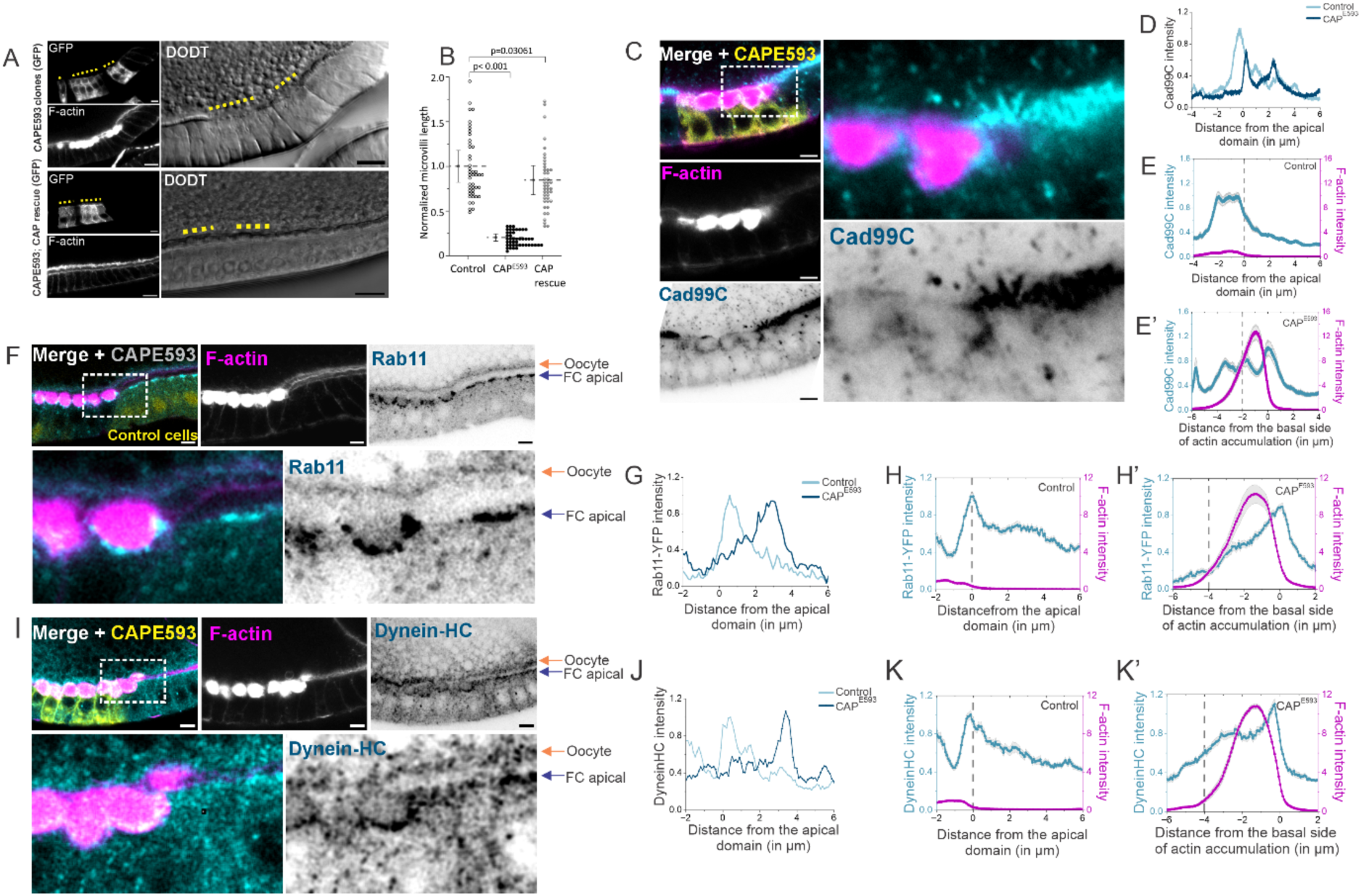
Apical actin accumulation halts Dynein and Rab11-mediated trafficking, leading to the retention of apical cargo Cad99C required for microvilli biogenesis. **(A) Loss of apical microvilli in CAP mutant cells.** Mosaic follicular epithelium with CAP mutant cells (GFP positive, dashed line). The gradient contrast (DODT) image shows the loss of apical microvilli in CAP mutant cells (dashed line). CAP mutant cells rescued with wild-type CAP expression (CAP rescue, GFP positive, dashed line) show the rescue of F-actin accumulation and apical microvilli. Scale bar = 10 µm. **(B) Quantification of microvilli length relative to control.** Quantification of microvilli length normalized with the control. The bar shows the ± 0.5 SD, and the dotted line represents the mean value. One-way ANOVA with post-hoc test was used to determine the statistical significance between different groups. Control = 45 cells, CAPE593 = 45 cells; CAP rescue = 43 cells. N = 9, 11, 7 egg chambers. **(C) Microvilli-specific atypical Cad99C localises beneath accumulated actin in CAP mutant cells**. Mosaic follicular epithelium with CAP mutant cells (GFP, pseudo-colored yellow, dashed line) stained for F-actin (phalloidin, magenta) and immunostaining against microvilli-specific atypical Cadherin, Cad99C (cyan). Zoomed-in inset (dotted box) highlights the retention of Cad99C (cyan), below the accumulated F-actin (magenta) and failure to reach the apical side of the follicle cells. Scale bar = 10 µm. **(D) Line scan intensity profile of Cad99C in control and CAP mutant cells**. The intensity profile of Cad99C is plotted for representative control (light cyan) and CAP mutant (dark cyan) cells, revealing a shift in Cad99C localisation at the apical region of CAP mutant cells**. (E) Comparative line scan of filamentous actin and Cad99C.** Mean line scan intensity profiles of F-actin (magenta) and Cad99C (cyan) for (E) control and (E’) CAP mutant follicle cells. In (E’), the x-axis represents the distance from the basal side of the accumulated F-actin to normalise for the variation in the space occupied by the accumulated actin in the mutant. The dotted line marks the apical-most boundary of the follicle cells, just below the microvilli (which in control cells are Cad99C positive). Data are presented as mean ± SEM; n = 15 cells, N = 6 egg chambers. **(F) Rab11-positive endosomes localise beneath apical actin in CAP mutant cells**. Mosaic follicular epithelium with CAP mutant cells marked by the absence of nuclear RFP (nlsRFP negative, pseudo-colored yellow), expressing YFP-tagged Rab11(pseudo colored cyan), stained with phalloidin against F-actin. Zoomed-in inset (dotted box) shows that the apically accumulated F-actin halts the apical trafficking of Rab11 endosomes. The orange arrow highlights the Rab11-YFP localisation on the oocyte cortex, and the blue arrow highlights the localisation on the apical surface of the follicle cells (FC). Scale bar = 10 µm. **(G) Line scan intensity profile of Rab11 in control and CAP mutant cells**. The intensity profile of Rab11-YFP is plotted for representative control (light cyan) and CAP mutant (dark cyan) cells, revealing a shift in Rab11 localisation at the apical region of CAP mutant cells. **(H) Comparative line scan of filamentous actin and Rab11.** Combined line scan intensity profiles of F-actin (magenta) and Rab11-YFP (cyan) for (H) control and (H’) CAP mutant follicle cells. In (H’), the x-axis represents the distance from the basal side of the accumulated F-actin to normalize for the variation in the space occupied by the accumulated actin in the mutant cells. The dotted line marks the apical boundary of the follicle cells, just below the microvilli. Data are presented as mean ± SEM; n = 20 cells, N = 6 egg chambers. **(I) Dynein accumulates basally to the actin accumulation.** Mosaic follicular epithelium with CAP mutant cells (GFP positive, pseudocoloured yellow) stained for F-actin (magenta) with phalloidin and immunostained against motor protein dynein heavy chain (Dynein-HC, pseudocoloured cyan). Zoomed-in inset (dotted box) shows the accumulation of Dynein beneath the apically accumulated F-actin. The orange arrow highlights the Dynein-HC localisation on the oocyte cortex, and the blue arrow highlights the Dynein-HC localisation on the apical surface of the follicle cells (FC). Scale bar = 10 µm. **(J) Line scan intensity profile of dynein in control and CAP mutant cells.** The intensity profile of Dynein heavy chain is plotted for representative control (light cyan) and CAP mutant (dark cyan) cells, revealing a shift in Dynein localisation at the apical region of CAP mutant cells. **(K) Comparative line scan of filamentous actin and dynein.** Mean line scan intensity profiles of F-actin (magenta) and Dynein (cyan) are presented for (K) control and (K’) CAP mutant follicle cells. In (K’), the x-axis represents the distance from the basal side of the accumulated F-actin to normalise for the variation in the space occupied by the accumulated actin in the cells. The dotted line marks the apical boundary of the follicle cells, just below the actin-rich microvilli. Data are presented as mean ± SEM; n = 11 cells, N = 4 egg chambers.

### Excess apical actin impairs microtubule organization in epithelial cells *in vivo*

During initial characterization of CAP mutants Baum & Perrimon, 2001 did not detect obvious defects in microtubules. However, experiments *in vitro* and in cultured cells have revealed that organization of actin filaments can prevent or guide the growth of microtubules (Colin et al., 2018; Gélin et al., 2023; Gupton & Waterman-Storer, 2006). Therefore, we decided to test this possibility *in vivo* in CAP mutant epithelial cells. To achieve a pronounced accumulation of apical actin, we used longer recovery time after generation of mutant clones compared to that used by Baum and coworkers. We found that, the apical region of CAP mutant cells, characterized by accumulated actin, exhibited decreased levels of stable acetylated microtubules (Figure 5 A-B’, Supl Figure 3A, A’) as well as α- and β-tubulin (Figure 5 C, D), indicating that microtubules were excluded from the region of actin accumulation. We also analyzed microtubule organization in stretch cells which are specialized flattened follicle cells. The flat morphology of stretch cells enables visualization of the cytoskeleton within the whole cell. In CAP mutant stretch cells, actin accumulation was predominant in one cytoplasmic region, which the stable acetylated microtubules were not able to penetrate (Supl Figure 3 B). Together, these findings reveal that dense actin structures exclude microtubules in epithelial cells *in vivo*.

**Figure 5.**
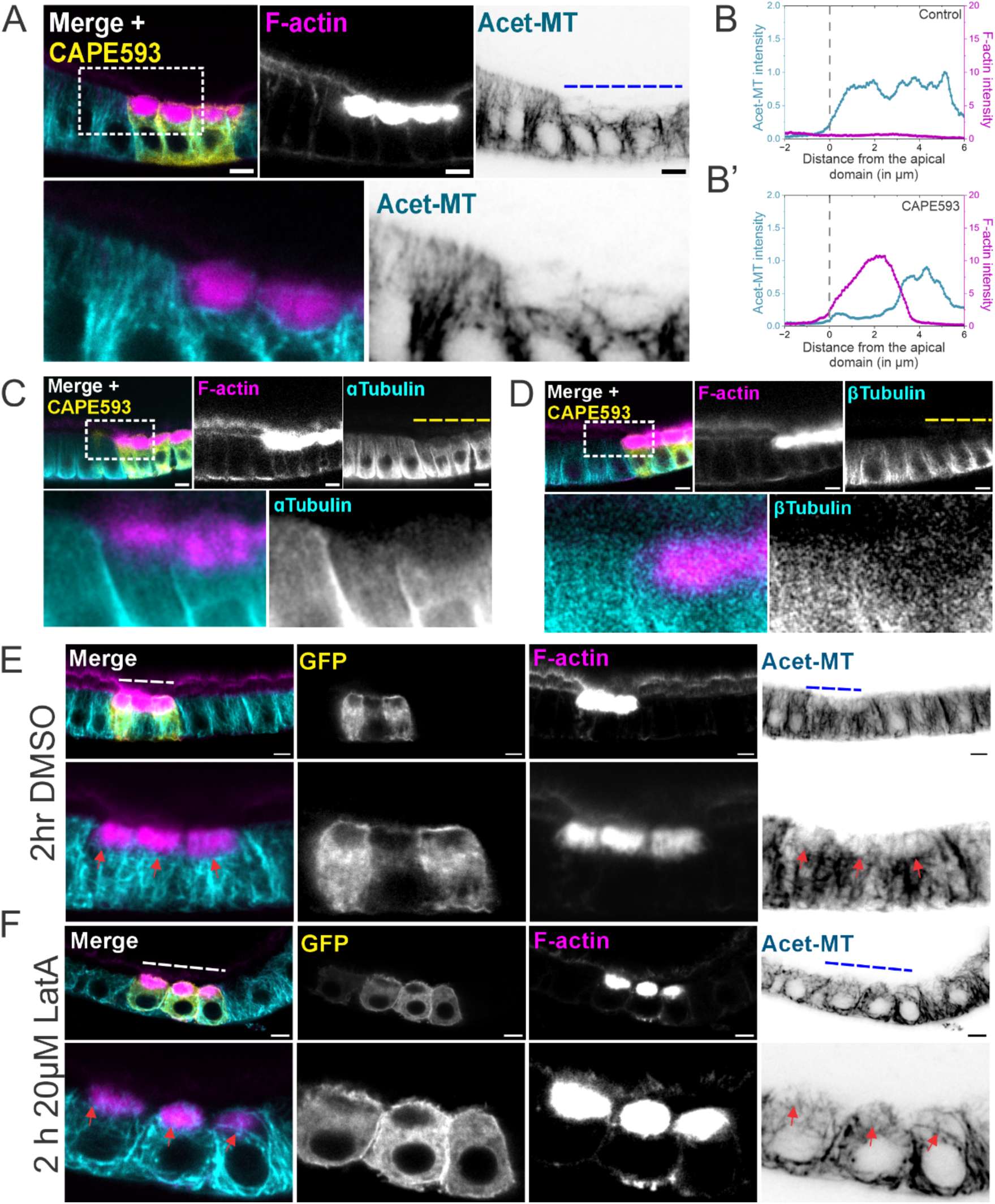
CAP loss of function results in F-actin accumulation and misorganized microtubules. **(A) Apical actin accumulation excludes acetylated microtubules in CAP mutant follicle cells.** A mosaic of follicular epithelium is shown, with CAP mutant cells highlighted by mCD8-GFP (GFP positive, pseudo-colored yellow, dashed line). The merged image illustrates the accumulation of filamentous actin at the apical region, as indicated by Phalloidin staining (pseudo-colored magenta). In contrast, acetylated microtubules (pseudo-coloured cyan) are absent from this region. Scale bar = 10 µm. Zoomed-in inset (dotted box) (Bertling et al., 2007)highlights the site of actin accumulation (magenta) devoid of acetylated microtubules. **(B) Line scan intensity profile** of the F-actin (magenta) and acetylated microtubules (cyan) for a single representative control (B) and CAP mutant follicle cell (B’). The dotted line marks the apical boundary of the follicle cells. **(C, D) Apical actin accumulation excludes microtubule α- and β-tubulin subunits in CAP mutant follicle cells** (GFP positive, pseudo-colored yellow, dashed lines). Zoomed-in inset (dotted box) shows apical region CAP mutant cells with actin accumulation (Phalloidin, pseudo-colored magenta) and the site of apical actin accumulation is devoid of (C) α-Tubulin (cyan), and (D) β-Tubulin (cyan). Scale bar = 10 µm. **(E, F) Latrunculin A treatment fails to resolve dense accumulated actin of CAP mutant cells but facilitates microtubule entry.** Mosaic follicular epithelium with CAP mutant cells marked with mCD8-GFP (GFP positive, pseudo-colored yellow, dashed line), stained for filamentous actin (stained with phalloidin, pseudo-colored magenta) and for acetylated microtubules (Acet-MT, pseudo-colored cyan), treated with (E) DMSO (control) and (F) 20 µM Latrunculin A for 2 hours. In (E) DMSO-treated egg chambers, the site of actin accumulation lacked the microtubules. (F) Upon treatment with Latrunculin A, acetylated microtubules are seen at the site of actin accumulation and membrane-bound mCD8-GFP is also localised at the site of actin accumulation, showing partial disassembly of the accumulated actin. Scale bar = 10 µm.

Unexpectedly, despite the remaining accumulation of filamentous actin, Lat A-treatment allowed some microtubules to pass through the accumulated actin (Figure 5 E, F). This result suggests that the accumulated actin undergo low levels of depolymerization, leading to a decrease in the total density following Lat A-treatment to allow microtubules to penetrate through the filamentous actin mass. Alternatively, a moderate reduction in the density of accumulated actin may have been caused by Latrunculin A itself, as it was reported to depolymerize actin filaments by severing and sequestering monomers (Fujiwara et al., 2018). Furthermore, the localization of membrane-bound mCD8-GFP was partially restored in the apical region of CAP mutant cells after Latrunculin A treatment (Figure 5 E, F). The result shows that microtubule-mediated transport of membrane-bound compartments to the apical regions is recovered by Latrunculin A treatment. Taken together, these results show that CAP loss of function leads to stable and dense apical actin structures, which likely forms a barrier that sterically prevents microtubule penetration and effective microtubule-based transport.

### Accumulation of apical actin affects localization of microtubule-actin crosslinking protein spectraplakin/Shot, involved in non-centrosomal microtubule organization

In many epithelial cells, microtubules are organized in a non-centrosomal array, with their minus-ends anchored at the apical cortex and their plus-ends extending basally (Bacallao et al., 1989; Bré et al., 1990; Clark et al., 1997). A central mechanism enabling this organization is the anchoring and stabilization of microtubule minus-ends at the apical cortex by Patronin, the *Drosophila* homolog of CAMSAP3, together with Shot (Short stop; *Drosophila* homolog of MACF1/ACF7 and DST/BPAG1), a spectraplakin that crosslinks microtubules to the actin cytoskeleton. Both Patronin and Shot show sharp apical localization in follicle cells (Khanal et al., 2016; Nashchekin et al., 2016).

Next, to determine how apical actin accumulation affects microtubule organization, we examined the localization of Patronin and Shot in CAP mutant follicle cells. We found that Patronin, a microtubule minus-end–stabilizing and anchoring protein, remained apically localized in mutant cells, although its intensity appeared increased compared to wild-type cells (Figure 6A–C’). In contrast, the microtubule–actin crosslinking protein Shot was enriched beneath the accumulated apical actin in CAP mutant cells, rather than showing the sharp apical localization observed in wild type (Figure 6D–F’). These results indicate that excess apical actin affects Patronin and Shot differently: whereas Patronin localization is largely maintained, Shot fails to localize properly to the apical cortex. Nevertheless, because Dynein, whose localization marks microtubule minus-ends, remain enriched in the apical region (Figure 4I–K), the overall apical–basal polarity of non-centrosomal microtubules in CAP mutant cells appears to be preserved.

**Figure 6.**
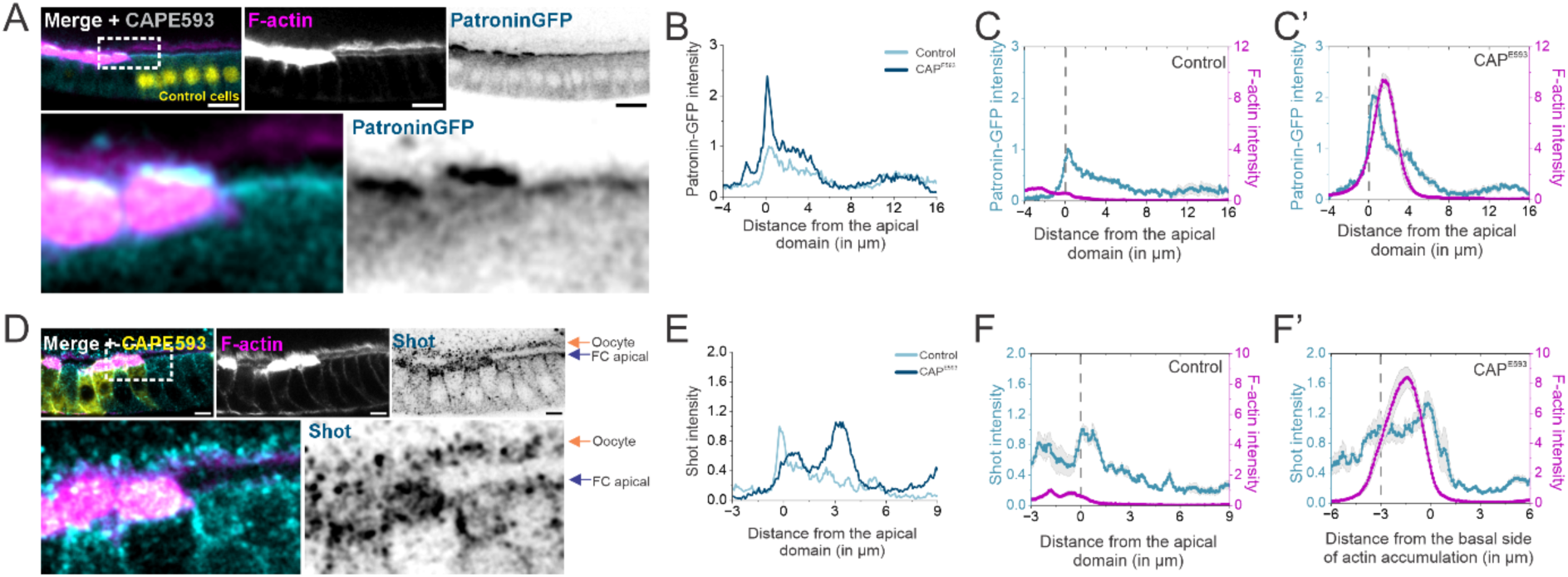
Localisation of microtubule minus-end anchoring proteins Patronin and actin-microtubule crosslinking protein Shot is impaired in CAP mutant cells. **(A) Apical localisation of microtubule minus-end anchoring protein Patronin.** Mosaic follicular epithelium with CAP mutant cells marked by the absence of nuclear RFP (nlsRFP (pseudocoloured yellow) negative) is shown. The merged image illustrates the accumulation of filamentous actin at the apical region, as indicated by Phalloidin staining (pseudo-colored magenta). Zoomed-in inset (dotted box), shows that the microtubule minus-end binding protein Patronin (Patronin-GFP, pseudo-colored cyan) is localized apically in both control cells (nlsRFP positive, pseudo-colored yellow) and CAP mutant cells (nlsRFP negative). However, the apical intensity of Patronin-GFP is higher in CAP mutant cells, and its localisation appears more diffuse. Scale bar = 10 µm. **(B) Line scan intensity profile of Patronin-GFP in control and CAP mutant cells.** The intensity profile of Patronin-GFP is plotted for representative control (light cyan) and CAP mutant (dark cyan) cells, revealing the increased localisation at the apical region of CAP mutant cells. **(C) Comparative line scan of filamentous actin and Patronin.** Mean line scan intensity profiles of F-actin (magenta) and Patronin-GFP (cyan) are presented for (C) control and (C’) CAP mutant follicle cells. The dotted line marks the apical boundary of the follicle cells, just below the actin-rich microvilli. Data are presented as mean ± SEM; n = 5 cells, N = 3 egg chambers. **(D) Impaired apical localisation of actin- microtubule crosslinking protein Shot.** Mosaic follicular epithelium with CAP mutant cells (GFP positive) staining F-actin (phalloidin) and antibody against Shot. The zoomed-in inset shows the disrupted apical localisation of actin-microtubule crosslinker Shot (pseudo-colored, cyan) at the site of actin accumulation. Shot is localised basal and around the accumulated actin. The orange arrow highlights the Shot localisation on the oocyte cortex, and the blue arrow highlights the localisation of Shot on the apical surface of the follicle cells (FC). Scale bar = 10 µm. **(E) Line scan intensity profile of Shot in control and CAP mutant cells**. The intensity profile of Shot is plotted for representative control (light cyan) and CAP mutant (dark cyan) cells, revealing a shift in Shot localisation at the apical region of CAP mutant cells. **(F) Comparative line scan of filamentous actin and Shot.** Combined line scan intensity profiles of F-actin (magenta) and Shot (cyan) for (F) control and (F’) CAP mutant follicle cells. In (F’), the x-axis represents the distance from the basal side of the accumulated F-actin to normalize for the variation in the space occupied by the accumulated actin in the mutant cells. The dotted line marks the apical boundary of the follicle cells, just below the microvilli. Data are presented as mean ± SEM; n = 5 cells, N = 3 egg chambers.

### Dense apical actin prevents normal microtubule organization and leads to nuclear mispositioning

Organization of the non-centrosomal microtubules is required for proper nuclear positioning in various cellular contexts such as in *Drosophila* follicular epithelium as well as in mammalian intestinal epithelial cells (Khanal et al., 2016; Muroyama et al., 2018; Nashchekin et al., 2016; Toya et al., 2016). Consequently, defects in nuclear positioning provide an independent readout to confirm microtubule-related phenotypes. We found that in CAP mutant cells, nuclei were mispositioned toward the apical region compared to the basal positions of the neighboring control cells (Figure 7 A). Surprisingly, reintroducing wild-type CAP into CAP mutant cells did not restore nuclear positioning, despite successfully rescuing actin accumulation, as well as earlier described phenotypes: organization of non-centrosomal microtubules, microvilli formation, and cytoplasmic mCD8-GFP localization in stage 10 (Figure 7 B, C). Quantitative analysis confirmed that the distance of the nuclei from the basal membrane was significantly greater in CAP mutants as compared to the controls (Figure 7 C). Furthermore, the nuclear mispositioning appeared even more evident in the rescued cells (Figure 7 B, C), possibly due to their better survival. Additionally, we did not observe any significant difference in the cell height between the control, CAP mutant and CAP wildtype rescue, suggesting that the observed nuclear position phenotype is independent of the cell height (Supl Fig 4 A). It is important to note that while we quantified nuclear positioning during stages 9-10, positioning occurs earlier, during stages 6-9 of oogenesis (Szikora et al., 2013). Thus, we asked if the CAP mutant phenotypes are visible at earlier stages of the follicle development. Indeed, the loss of CAP led to actin accumulation and disrupted apical microtubule organization in follicle cells during both early (stage 7, Figure 7 D) and late (stage 10, Figure 7 E) stages of oogenesis. However, the reintroduction of wild-type CAP only rescued actin accumulation and microtubule organization at stage 10, not at stage 7 (Figure 7 F, G). Further analysis showed that the CAP rescue construct was only minimally expressed during stages 6-7 (Supl Fig 4 B) indicating that the nuclear mispositioning phenotype, which develops during the early stages of oogenesis, could not be rescued by CAP expression at later stages. In conclusion, these results underscore the temporal requirement of CAP-mediated actin dynamics for establishing non-centrosomal microtubule arrays during early oogenesis for ensuring correct nuclear positioning.

**Figure 7.**
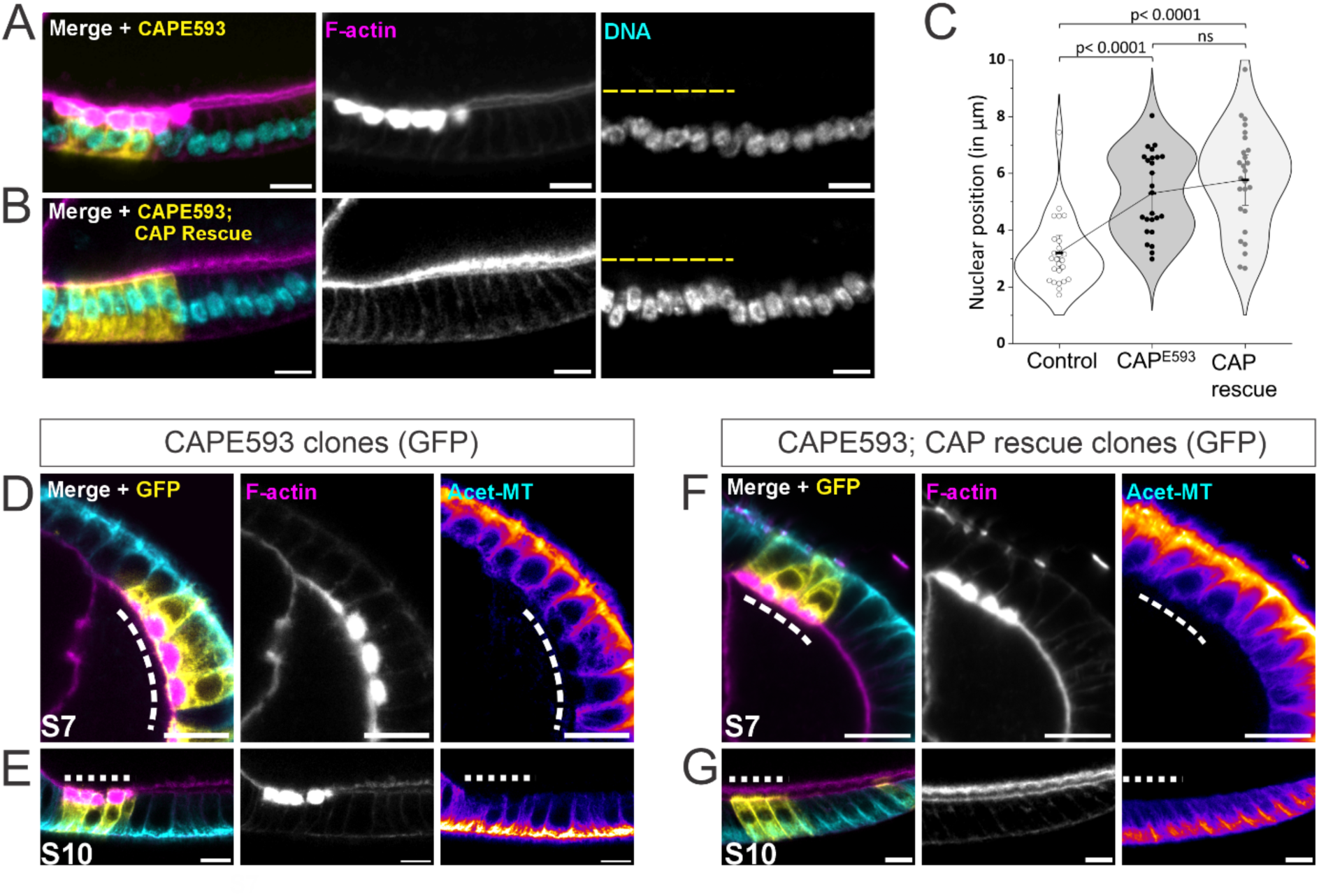
Apical actin accumulation during early oogenesis causes mispositioning of follicle cell nuclei. **(A) CAP mutant cell nuclei are mispositioned apically.** Mosaic follicular epithelium with CAP mutant cells marked with mCD8-GFP (GFP positive, pseudo-colored yellow, dashed line) and filamentous actin (stained with phalloidin, pseudo-colored magenta). In CAP mutant cells, nuclei (DNA stained with DAPI, pseudo-colored cyan) are positioned apically, while neighboring control cells (GFP negative) exhibit more basal nuclear positioning. Scale bar = 10 µm. **(B) Restoring CAP expression in CAP mutant cells rescues actin accumulation, but not the nuclear positioning phenotype.** Mosaic follicular epithelium with CAP mutant cells rescued by wild-type CAP expression (CAP rescue, GFP positive, pseudo-colored yellow, dashed line). While apical actin accumulation (filamentous actin stained with phalloidin, pseudo-colored magenta) is restored, the nuclei (DNA, cyan) remain mislocalized to the apical domain. Scale bar = 10 µm. **(C) Quantification of nuclear position in control, CAP mutant, and rescued cells.** A quantification graph is presented showing the nuclear position defined by the distance from the nucleus’s center to the cell’s basal side (in µm). The connecting line passes through the mean value of each group, and the whiskers represent the standard deviation. Kruskal–Wallis ANOVA followed by Dunn’s post hoc test was used to assess the statistical significance of the different groups. Control = 25 cells, CAPE593 = 25 cells, and CAP rescue = 26 cells. N = 6 egg chambers. **(D, E) CAP mutant cells display accumulated actin and microtubule disorganisation in both stage 7 and stage 10.** A cross-section of (D) stage 7 and (E) stage 10 mosaic follicular epithelium is shown, with CAP mutant cells highlighted by mCD8-GFP (GFP positive, dashed line). In both stages, CAP mutant cells display apical actin accumulation (filamentous actin stained with phalloidin, pseudo-colored magenta) and absence of acetylated microtubules (Acet-MT, pseudo-colored cyan). Scale bar = 10 µm. **(F) Actin accumulation and microtubule organization phenotypes are not rescued during early oogenesis.** A cross-section of stage 7 (s7) follicular epithelium, with CAP rescue clones highlighted by mCD8-GFP (GFP positive, pseudo-colored yellow, dashed line). The merged image demonstrates that apical actin accumulation is not rescued (filamentous actin stained with phalloidin, pseudo-colored magenta) nor is the organization of stable acetylated microtubules (Acet-MT, pseudo-colored cyan) in stage 7. Scale bar = 10 µm. **(G) Actin accumulation and microtubule organization phenotypes are successfully rescued in later stages of oogenesis.** A cross-section of stage 10 (s10) follicular epithelium is shown, with CAP wildtype rescue clones highlighted by mCD8-GFP (GFP positive, pseudo-colored yellow, dashed line). Both apical actin accumulation (filamentous actin stained with phalloidin, pseudo-colored magenta) and stable acetylated microtubules (Acet-MT, pseudo-colored cyan) are successfully restored. Scale bar = 10 µm.

## DISCUSSION

### Coordination of apical actin and microtubules

Our findings demonstrate that CAP-mediated apical actin turnover is needed for correct organization of non-centrosomal microtubule arrays in *Drosophila* follicular epithelium. The loss of function of Cyclase-Associated Protein (CAP, capt in *Drosophila*) leads to impenetrable ectopic apical actin forming a barrier that in turn disrupts apical organization of non-centrosomal microtubule arrays. As organization of noncentrosomal microtubules are disrupted, CAP mutant cells show defects in nuclear positioning, Dynein-based apical cargo trafficking, and the formation of apical microvilli. Our results highlight the role of CAP-mediated apical actin filament turnover in microtubule organization, and suggest that apical actin turnover plays an important role in cytoskeletal coordination of epithelial cells.

The coordination of apical actin and non-centrosomal microtubules has been earlier explored in the context of epithelial morphogenesis, focusing on the generation of actomyosin-mediated contractile and adhesive forces (Röper, 2020). Current literature describes microtubules as regulators of apical actomyosin pool and coordinating actomyosin-based forces during apical constriction (Booth et al., 2014b; Corrigall et al., 2007; Ko et al., 2019b; J.-Y. Lee & Harland, 2007). Additionally, both apical actin turnover and non-centrosomal microtubules have been recognized as important regulators of force balance in epithelial cells (Ikawa & Sugimura, 2018; Jodoin et al., 2015). How and whether actin turnover, microtubule organization and actomyosin-based contractility are co-orchestrated during epithelial morphogenesis requires further investigation.

It is well-known that actin filament architectures actively modulate microtubules *in vitro* and in cultured cells (Cheng et al., 2023; Gélin et al., 2023; Gupton & Waterman-Storer, 2006; Waterman-Storer & Salmon, 1997). Also, loss of profilin, actin-binding protein facilitating the assembly of actin monomers into pre-existing actin filaments, results in changes in microtubule organization in cells, and therefore has been suggested to regulate both actin and microtubules (Henty-Ridilla et al., 2017; Liu et al., 2022; Nejedla et al., 2016). However, it has been recently postulated that the microtubule phenotypes associated with profilin loss are likely to result from adaptive responses of microtubules to changes in the actin cytoskeleton architecture and not as direct effect of loss of the profilin (Cisterna et al., 2024).

One mechanism by which filamentous actin directly regulates microtubules is through steric exclusion, where dense actin architectures can obstruct microtubule plus-end growth or cause their disassembly (Colin et al., 2018). Likely, the observed dense and persistent apical actin acts as a physical barrier to microtubules in CAP mutant cells. Thus, by promoting actin filament turnover CAP regulates actin filament architecture and density and may indirectly affect microtubules organization. This is supported by the observation that prolonged Latrunculin A treatment “loosens” the accumulated actin in CAP mutant cells, allowing microtubules to pass through. The actin cytoskeleton is known to sterically prevent microtubules in sites where the two cytoskeletal components converge in crowded spaces, such as at the leading edge of migrating cells or in neural growth cones (Dogterom & Koenderink, 2018; Pimm & Henty-Ridilla, 2021). In addition to regulating microtubules at the cell periphery, dense actin filaments in the centrosome reduce the number of centrosomal microtubules (Farina et al., 2019; Inoue et al., 2019). This suggests that dynamic changes in actin filament densities can be regulated according to the needs of microtubule assembly.

### Dense apical actin prevents normal organization on non-centrosomal microtubules and results in nuclear mispostioning

The nuclear positioning phenotype of CAP-mutant cells suggests that apical actin turnover during early oogenesis is important for organization of non-centrosomal microtubules. In actively dividing cells, microtubules are typically nucleated by the centrosome, whereas in postmitotic epithelial cells, microtubules are organized into non-centrosomal arrays (Muroyama & Lechler, 2017). The transition from centrosomal to non-centrosomal microtubules coincides with the termination of cell divisions. In the follicular epithelium, cells divide until stage 6 (Klusza & Deng, 2011), suggesting that transition to non-centrosomal microtubules occurs thereafter. In follicle cells, nuclear positioning appears to be temporally connected to the establishment of non-centrosomal microtubule arrays during stages 6-9. In CAP mutant cells with stabilized apical actin, nuclei are positioned more apically compared to wild-type cells. Correct nuclear positioning in follicle cells is determined by the growing plus-ends of apically anchored microtubules that exert force to push nuclei to the correct apical-basal position (Szikora et al., 2013). Thus, the nuclear positioning phenotype in CAP mutant cells is likely caused by disorganization of non-centrosomal microtubules. These findings suggest that apical actin turnover is critical for correct nuclear positioning at the time when non-centrosomal microtubule arrays are established.

Correct organization of non-centrosomal microtubules is dependent on cortical anchoring of microtubule-minus ends mediated by Patronin and Shot. In addition, both Patronin and Shot are required for ensuring correct nuclear positioning across various cell types, from the *Drosophila* follicular epithelium to mammalian intestinal cells (Khanal et al., 2016; Muroyama et al., 2018; Nashchekin et al., 2016; Toya et al., 2016). Normally, Shot and Patronin exhibit sharp apical localization. In CAP mutant cells, however, Shot is localized ectopically beneath the accumulated actin. Ectopic Shot localization in CAP mutant cells may be explained by the fact that Shot’s ability to bind microtubule lattices and promote actin-microtubule crosslinking is enhanced by increased filamentous actin densities (Applewhite et al., 2010, 2013; Takács et al., 2017). It is thus possible that in CAP mutant cells, accumulated actin traps Shot to ectopic sites, where Shot crosslinks microtubules and actin further impairing cytoskeletal coordination. To conclude, we demonstrate that efficient apical actin turnover mediated by CAP is needed for correct organization of non-centrosomal microtubule arrays to promote correct nuclear positioning.

### Relevance of our findings and similarities to other polarized cells

Our current study enhances the understanding of CAP-mediated actin turnover in non-centrosomal microtubule organization. One of the earliest phenotypes associated with CAP loss-of-function was a defect in *Drosophila* oocyte polarity (Baum et al., 2000). In *Drosophila*, oocyte polarity is established by non-centrosomal microtubule arrays, that transport polarity-determining mRNAs to the anterior and posterior ends (St Johnston, 2023). In CAP mutant oocytes, these mRNAs are mislocalized, suggesting a defect in microtubule polarity. However, the mechanism by which CAP contributes to microtubule-related phenotypes has been incompletely understood. Like follicle cells, in oocytes transition from centrosomal to non-centrosomal microtubule array establishment is critical for oocyte polarity. In CAP mutant oocytes, excessive actin accumulates at the anterior end (Baum et al., 2000). Intriguingly, this is the cortical site where non-centrosomal microtubule arrays are normally anchored via Shot and Patronin (Nashchekin et al., 2016). Our results show that in follicle cells CAP-mediated actin turnover is critical for proper localization of Shot. Whether CAP-mediated actin turnover is also required for the microtubule minus targeting in oocytes remains to be investigated.

CAP is well-established for promoting actin dynamics in heart and skeletal muscle cells, as well as in neuronal cells. Its loss results in actin accumulation, which impairs muscle development and contraction, and affects neuronal growth cones and dendritic spines (Field et al., 2015; Heinze et al., 2022; Kepser et al., 2019; Kumar et al., 2016; Peche et al., 2013; Schneider et al., 2021). Furthermore, the loss of CAP activity, along with excessive actin accumulation, has been linked to diseases such as nemaline myopathy and Alzheimer’s disease (Gurunathan et al., 2022; Pelucchi et al., 2020). Our results shows that disruption of actin turnover and excessive accumulation of actin in CAP mutant follicle cells can disrupt microtubule organization and microtubule-based transport. Interestingly, in both muscle and neuronal cells, microtubules are organized in a non-centrosomal manner (Bugnard et al., 2005; Kapitein & Hoogenraad, 2015; Stiess et al., 2010; Tassin et al., 1985). However, the impact of actin accumulation on microtubules has not been extensively studied muscle cells. In neuronal cells it is known that persistent cofilin-actin rods disrupt microtubules and impair microtubule-based trafficking (Cichon et al., 2012; Medina et al., 2008; Minamide et al., 2000). Therefore, future studies are needed to determine how disruption of actin turnover affects non-centrosomal microtubule arrays in other cellular contexts, further elucidating the intricate interplay between actin dynamics and microtubule organization in cells.

## MATERIALS AND METHODS

### Drosophila stocks

All the fly lines used in this study were obtained from Bloomington *Drosophila* Stock Center or shared as gifts from other research groups. *Drosophila* fly line CaptE539, an allele of Act up or Capulet (described as CAP) encoding a nonsense mutation at tryptophan 141 (described in (Benlali et al., 2000), BDSC 5944) was used to generate CAP (Cyclase-associated protein) chimeric loss-of-function mutant clones. Mitotic mutant clones in the follicular epithelium were generated using FLP/FRT systems, hsFLP; TubGal80, FRT40A; TubP-Gal4, UAS-mCD8-GFP/ TM6B, Tb (MARCM 40, Gifted by Pernille Rørth) and hsflp; UbimRFPnls FRT40/cyo. Used genetypes for experiments are described in Table 2 and Supplementary Table 1.

**Table 2:** Drosophila genotypes used.

| Figure | Panel | Genotype | Abbreviation | Source |
| --- | --- | --- | --- | --- |
| 1 | B | w1118 | w1118 |  |
|  | C-E | yw,hsFLP/w;TubGal80,FRT40A/Capt<sup>E593</sup><br>FRT40A; TubP-GAL4, UAS-mCD8-GFP / + | CAP<sup>E593</sup> | BDSC 5944 |
| 2 | B | yw,hsFLP/w;TubGal80,FRT40A/Capt<sup>E593</sup><br>FRT40A; TubP-GAL4, UAS-mCD8-GFP / + | CAP<sup>E593</sup> | BDSC 5944 |
|  | C | yw,hsFLP/w;TubGal80,FRT40A/Capt<sup>E593</sup><br>FRT40A; TubP-GAL4, UAS-Capt/ UAS-mCD8-GFP | CAP<sup>E593</sup>; CAP<br>Rescue | BDSC 5944<br>UAS-Capt, in<br>this ms |
|  | D | yw,hsFLP/w;TubGal80,FRT40A/Capt<sup>E593</sup><br>FRT40A; TubP-GAL4, UAS-Cap<sup>tm6</sup>/ UAS-mCD8-GFP | CAP<sup>E593</sup>;<br>CAP<sup>tm6</sup>mutant | BDSC 5944<br>UAS-Cap<sup>tm6</sup>,<br>in this ms |
|  | E | yw,hsFLP/w;TubGal80,FRT40A/Capt<sup>E593</sup><br>FRT40A; TubP-GAL4, UAS-Chic/ UAS-mCD8-GFP | CAP<sup>E593</sup>;<br>ProfilinOE | BDSC 5944<br>Lynn Cooley |
| 3 | A, C,<br>D, E | w,hsFLP/w; TubGal80, FRT40A/Capt<sup>E593</sup><br>FRT40A; TubP-GAL4, UAS-mCD8-GFP / + | CAP<sup>E593</sup> | BDSC 5944 |
|  | B | yw,hsFLP/w;TubGal80,FRT40A/Capt<sup>E593</sup><br>FRT40A; TubP-GAL4, UAS-Capt/ UAS-mCD8-GFP | CAP<sup>E593</sup>; CAP<br>Rescue | UAS-Capt, in<br>this ms |
| 4 | A, C, I | w,hsFLP/w; TubGal80, FRT40A/Capt<sup>E593</sup><br>FRT40A; TubP-GAL4, UAS-mCD8-GFP / + | CAP<sup>E593</sup> | BDSC 5944 |
|  | F | hsflp; UbimRFPnls, FRT40/Capt<sup>E593</sup><br>FRT40A; Rab11EYFP/+ | CAP<sup>E593</sup>;<br>Rab11YFP | BDSC 5944<br>BDSC 62549 |
|  | A | yw,hsFLP/w;TubGal80,FRT40A/Capt<sup>E593</sup><br>FRT40A; TubP-GAL4, UAS-Capt/ UAS-mCD8-GFP | CAP<sup>E593</sup>; CAP<br>Rescue | BDSC 5944<br>UAS-Capt, in<br>this ms |
| 5 | A, C-F | w,hsFLP/w; TubGal80, FRT40A/Capt<sup>E593</sup><br>FRT40A; TubP-GAL4, UAS-mCD8-GFP / + | CAP<sup>E593</sup> | BDSC 5944 |
| 6 | A | hsflp;UbimRFPnls,FRT40/Capt<sup>E593</sup><br>FRT40A; ubi.Patronin-GFP/+ | CAP<sup>E593</sup>;<br>Patronin-GFP | BDSC 5944<br>BDSC 55129 |
|  | B | yw,hsFLP/w;TubGal80,FRT40A/Capt<sup>E593</sup><br>FRT40A; TubP-GAL4, UAS-mCD8-GFP / + | CAP<sup>E593</sup> | BDSC 5944 |
| 7 | A, D,<br>E | w,hsFLP/w;TubGal80,FRT40A/Capt<sup>E593</sup><br>FRT40A; TubP-GAL4, UAS-mCD8-GFP / + | CAP<sup>E593</sup> | BDSC 5944 |
|  | B, F,<br>G | yw,hsFLP/w;TubGal80,FRT40A/Capt<sup>E593</sup><br>FRT40A; TubP-GAL4, UAS-Capt/ UAS-mCD8-GFP | CAP<sup>E593</sup>; CAP<br>Rescue | BDSC 5944<br>UAS-Capt, in<br>this ms. |

CAP loss of function clones were rescued with UAS fly lines encoding for wildtype CAP and specific point mutations that abolished different biochemical activities of CAP. These UAS-CAP fly lines were generated in-house for this project (see Table 1). UAS-Chic, encoding for profilin was kind gift from Lynn Cooley. Endosomal trafficking in mutant clones was analysed by combining Rab11-EYFP (BDSC 62549) with CaptE593. Localization of F-actin crosslinker α-actinin is visualized by combining CaptE593 with α-actinin-GFP (BDSC 602268). Localization of AIP1 was analyzed using AIP1- protein trap AIP1GFP (BDSC 50824). Combining CaptE593 and Ubi-p63E.Patronin GFP (BDSC 55129) were used to analyze the localization of microtubule minus-end binding protein Ppatronin. All fly stocks were maintained at 18°C and crossed at 25°C in a standard food medium.

### Generation of CAP domain mutants

Site-directed mutagenesis of selected amino acids to alanins (Table 1) was performed within the longer cDNA of CAPT-RB (RH0878) isoform. The parts corresponding to CAPT-RA isoform from the mutated CAPT-RB (RH0878, FlyBase ID FBtr0100023) isoform were cloned into the pUAST-attB vector. PCR was amplified using DNA polymerase enzyme KAPA HiFi (ReadyMix x 2, #KK2601, KAPA Biosystems; 30-60 sec/kb) following the recommended protocol. Resulting plasmids were verified with sequencing. The transgenic flies were generated by BestGene inc. *Drosophila* embryo injection services.

### Generation of mutant clones

For the generation of mutant clones, 2–4-day-old female flies were transferred to empty food vials and given two heat shocks at 37°C for one hour with a 5-hour interval between each heat shock. After the two heat shocks, flies were kept in dry yeast and allowed to recover for 3 days at 25°C. For rescue experiments clones were generated up to 9 days prior dissections, in order toto ensure the disappearance of endogenous CAP. After the recovery ovaries were dissected from these flies. The flies were transferred to a fresh food medium containing wet yeast overnight before the dissection.

### Producing recombinant Capt protein

D. melanogaster Capt protein was expressed using pCoofy18 expression plasmid (a kind gift from Sabine Suppmann, Addgene plasmid #43975) in BL21(DE3) E. coli (Sigma-Aldrich). Protein purification was carried out essentially as described in Kotila et al. 2018. In brief, collected bacterial cells were lysed in 50 mM Tris-HCl, 150 mM NaCl, 25 mM imidazole, pH 7.5 buffer containing protease inhibitors. Supernatant was clarified by centrifugation and loaded to HisTrap FF crude prepacked column (GE Healthcare). Bound protein was eluted with imidazole gradient. Fractions containing Capt were pooled and dialyzed O/N at 4°C in the presence of 3C protease in 50 mM Tris-HCl, 150 mM NaCl, 1 mM DTT, pH 8.0 to remove 10xHis fusion tag. Dialyzed Capt protein was concentrated with Amicon Ultra-4 centrifugal filter (Merck) and loaded to HiLoad 16/600 SD 200 pg (GE Healthcare) column equilibrated in 5 mM HEPES, 50 mM NaCl, 0.2 mM DTT, 0.01% NaN3, pH 8.0 buffer. Fractions containing Capt protein were concentrated as described above and snap frozen at 5-10 mg/ml concentration using liquid N2 and stored at -80°C.

### Generation of Capt antibody

Ig fractions of Capt-immunized rabbit serum (Pineda Antibody Service) were purified by affinity chromatography from resin-antigen-coated beads. 500 µL of fraction was eluted from the beads with 5 mL 0.1 M glycine-HCl (pH 2.8). The specificity of the collected fractions was determined with the western blot of fly lysates of wild type (W1118) and captE593 lysates. Additionally, immunostaining with purified anti-Capt antibody showed decreased intensity in CaptE593 clones relative to the control cells.

### Dissection and Immunostaining

Ovaries were dissected in ice-cold PBS and fixed in 4% paraformaldehyde for 20 min. For acetylated α-tubulin staining, the ovaries were dissected in room temperature PBS and fixed in 8% paraformaldehyde for 10 min. For Dynein heavy chain (Dynein-HC) and Shot staining, ovaries were dissected in cold PBS and fixed for 10 min using cytoskeleton buffer without glutaraldehyde, with 10% sucrose, 8% paraformaldehyde and 0.05% Triton-X, thereafter ovaries were washed once with PBS and quenched with sodium borohydride (NaBH4) for 10 min. The egg chambers are permeabilized in 0.1% BSA in 0.5% PBST and blocked with blocking buffer 2.5% BSA in 0.5% PBST. The primary and secondary antibodies were diluted in the blocking buffer, and the samples were incubated overnight at 4°C and for 2 hours at room temperature respectively.

Primary antibodies used were rabbit anti-Capt (1:400, generated in the lab), mouse anti-acetylated αtubulin (1:100, Santa Cruz biotechnology 6-11B-1), rabbit anti-αTubulin (1:50, Abcam Ab18251), mouse anti-βTubulin (1:50, DSHB E7), mouse anti-Shot (1:25, DSHB mABRod1), mouse anti-Cnx99A (1:25, DSHB Cnx99A 6-2-1), mouse anti-Golgin84 (1:25, DSHB Golgin84 12-1), mouse anti-DyneinHC (1:25, DSHB 2C11-2), rabbit anti-Cad99C (1:2000, gift from Christian Dahmann lab; Schlichting et al., 2005), mouse anti-profilin (1:25, DSHB chi 1J), Twinfilin (gift from Tapio Heino; Wahlström et al., 2001), mouse anti-Ena (1:25, DSHB 5G2), rat anti-Tropomyosin (1:25, DSHB BB5/37.1) and rabbit anti-Phospho-Myosin II (1:300, Cell Signalling Technologies). Secondary antibodies, donkey anti-mouse Alexa 647 and goat anti-rabbit Alexa 647 (from Thermofisher Scientific), were used in a 1:300 dilution. 1µg/mL DAPI and Phalloidin (1:300, from Thermofisher Scientific) for staining the nucleus and F-actin, respectively.

### Transmission electron microscopy

Ovaries were dissected in cold 0.1M phosphate buffer (pH 7.4) and fixed in 2.5% glutaraldehyde (EM-grade) and 2% paraformaldehyde in 0.1M phosphate buffer for 1 hour at room temperature and then continued at 4°C overnight. Egg chambers were then washed with cold phosphate buffer on ice and post-fixed with 1% osmium tetroxide in phosphate buffer for 1 hour on ice, later washed with dH2O and stained with 1% uranyl acetate in dH2O at 4°C in the dark for 1 hour. After post-fixation and EMem block staining, egg chambers were dehydrated through a gradient of ethanol, rinsed with propylene oxide and gradually embedded into TAAB Hard epoxy resin. 60-nm thin sections were cut using an ultramicrotome (Leica EM Ultracut UC7, Leica Mikrosysteme GmbH, Austria), collected on Pioloform-coated single slot Copper grids and poststained with uranyl acetate and lead citrate. TEM imaging was done using JEM-1400 (JEOL Ltd, Japan) equipped with an Orius SC 1000B bottom-mounted CCD-camera (Gatan Inc., USA) at accelerating voltage of 80 kV and magnifications ranging from 8000 × to 100,000 ×. During TEM imaging the CAP mutant cells in chimeric tissue were identified based on the phenotypes correlating with the immunostaining such as loss of ER and golgi as well as reduced microvilli and perivitelline space.

### Confocal microscopy and image analysis

Confocal images were acquired using Leica SP8 Upright microscope using 20x and 63x oil immersion objectives. Image processing and analysis were conducted using ImageJ software. Fluorescence intensity in the apical region was calculated as the average value of all pixels within the ROI spanning the apical subcellular region. Data for the Line scan intensity plots for CAPE593 were acquired from line ROIs parallel to the apicobasal axis from the cross-section of the follicle cells. The distance is adjusted such that 0 in the x-axis of the graphs corresponds to 25% of the total F-actin intensity. The signal intensities are normalised to the maximum of the control sample, and maximum-minimum normalisation was performed for the mean value for plotting. For quantifying the nuclear position, the distance between the center of the nucleus to the basal surface (in µm) is measured as a proxy to show the shift in the position of the nucleus.

### Statistical analysis

The outcomes were derived from a minimum of two autonomous experiments, employing multiple egg chambers for analysis with reproducibility. Before performing the statistical analysis normality of the data set was analyzed using the W test or the Shapiro-Wilk test. For the normally distributed data sets, parametric tests, paired t-tests were used and for non-parametric data sets, Mann-Whitney tests were used to assess the significance of differences between the control and the CAPE593. For data with more than two groups Kruskal-Wallis H test or one-way ANOVA was conducted. The analysis was performed using OriginPro 2025 software.

## Supporting information

Babu et al supplementary material

## ACKNOWLEDGEMENTS

We thank Lappalainen lab for helpful discussions during this work, and V. Hietakangas and P. Lappalainen for critical reading providing feedback on the manuscript. We thank A. Aalto, A. Haapalainen, J. Koivula, A. Poukova, P. Puonti, S. Rytövuori, and M. Tolonen for excellent technical assistance. We thank Bloomington *Drosophila* Stock Center, V. Hietakangas, P. Rorth, F. Besse and L. Cooley for providing fly stocks. We thank Developmental studies Hybridoma bank, C. Dahmann and T. Heino for providing antibodies. This study was facilitated by the University of Helsinki *Drosophila* core facility (Hi-Fly) and Institute of Biotechnology Light Microscopy Unit (LMU), Electron Microscopy Unit (EMBI), and Biomedicum Imaging Unit (BIU) of the University of Helsinki, supported by HiLIFE and Biocenter Finland.

## Author contributions

M.P. conceptualization. M.P. and A.B. experimental design. A.B. carried out majority of the experiments, validated and analyzed the data. S.M made some of the initial discoveries and performed the CAP mutant rescue experiments. K.K. and T.K. contributed to cloning and protein purification for antibody generation. V.H. and J.M. provided reagents and expertise. V.H., J.M. and M.P Supervision. A.B. and M.P. wrote the manuscript with input from all authors.

