## Supplementary material for "Apical actin filament turnover mediated by cyclase-associated protein is required for organization of non-centrosomal microtubules in epithelium": Babu et al supplementary material

**Babu et al., Supplementary figures**

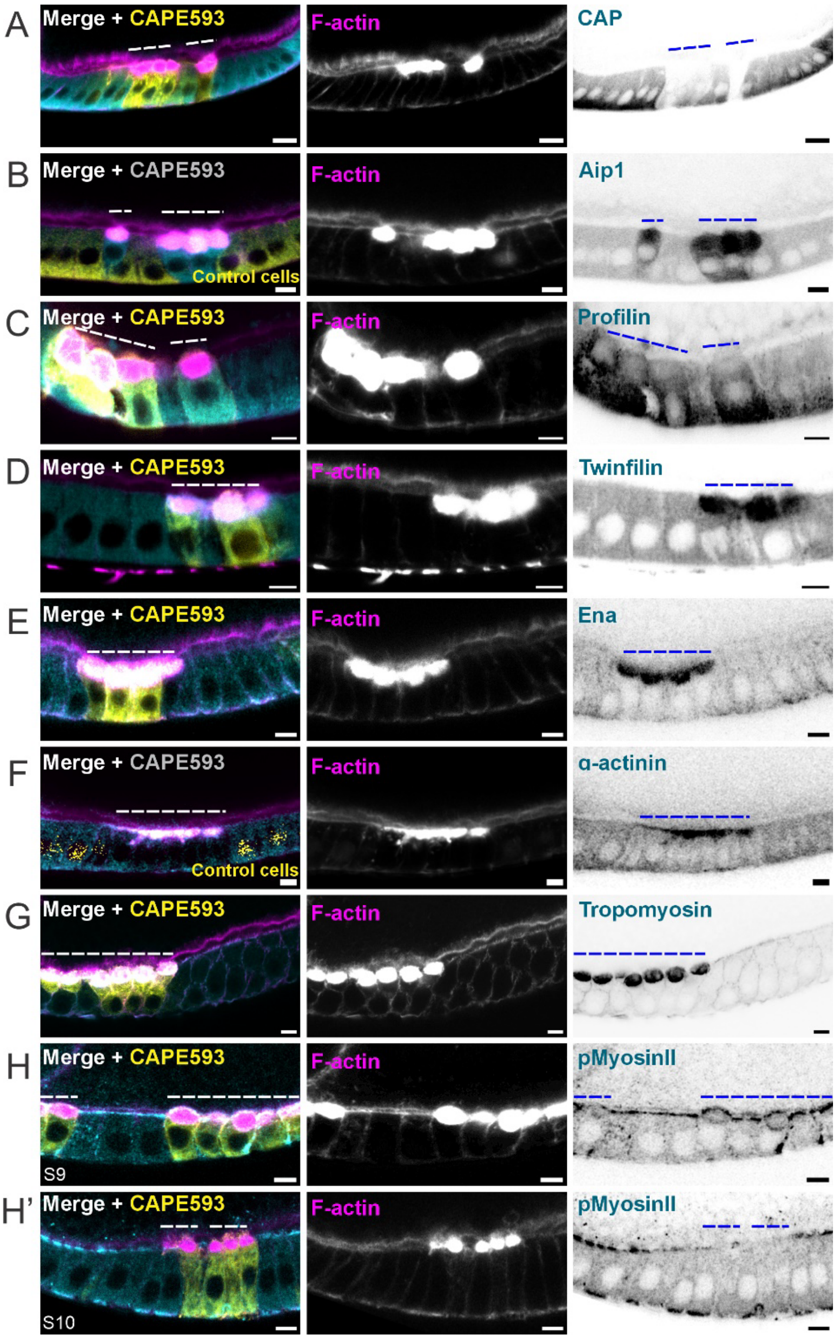


**Supplementary Figure 1. Localization of actin-binding proteins in CAP mutant cells at the site of apically accumulated F-actin** Mosaic follicular epithelium is shown, with CAP mutant cells highlighted with dashed line. The merged image illustrates the accumulation of filamentous actin at the apical region, indicated by Phalloidin staining (pseudo-colored magenta). **(A) CAP** is normally diffusely localized in cytoplasm (detected by immunostaining, pseudo-colored cyan) and is lost in CAP mutant cells (GFP positive, pseudo-colored yellow, dashed line). **(B) Actin Interacting Protein 1** (AIP1, GFP-tagged, pseudo-colored cyan) levels are increased in CAP mutant cells and it is enriched at the site of actin accumulation. **(C) Profilin,** promoting actin polymerization (detected by immunostaining, pseudo-colored cyan) levels are increased basally in CAP mutant cells, but decreased at the site of apical F-actin accumulation. **(D) Twinfilin** (detected by immunostaining, pseudo-colored cyan) is enriched at the site of actin accumulation in CAP mutant cells. **(E) Ena** (detected by immunostaining, pseudo-colored cyan) is enriched at the site of actin accumulation in CAP mutant cells. **(F) α-actinin** (pseudo-colored cyan) is enriched at the site of actin accumulation in CAP mutant cells (nlsRFP negative, dashed line). **(G) Tropomyosin** (detected by immunostaining pseudo-colored cyan) is enriched at the site of actin accumulation in CAP mutant cells. **(H, H') Active non-muscle myosin II** localization in CAP mutant cells depends on the stage. (**H**) **In stage 9** cells, non-muscle myosin II is localised below the accumulated F-actin, while (H') **in stage 10** cells, myosin II is absent from the apical domain detected by immunostaining against phospho-myosin II.

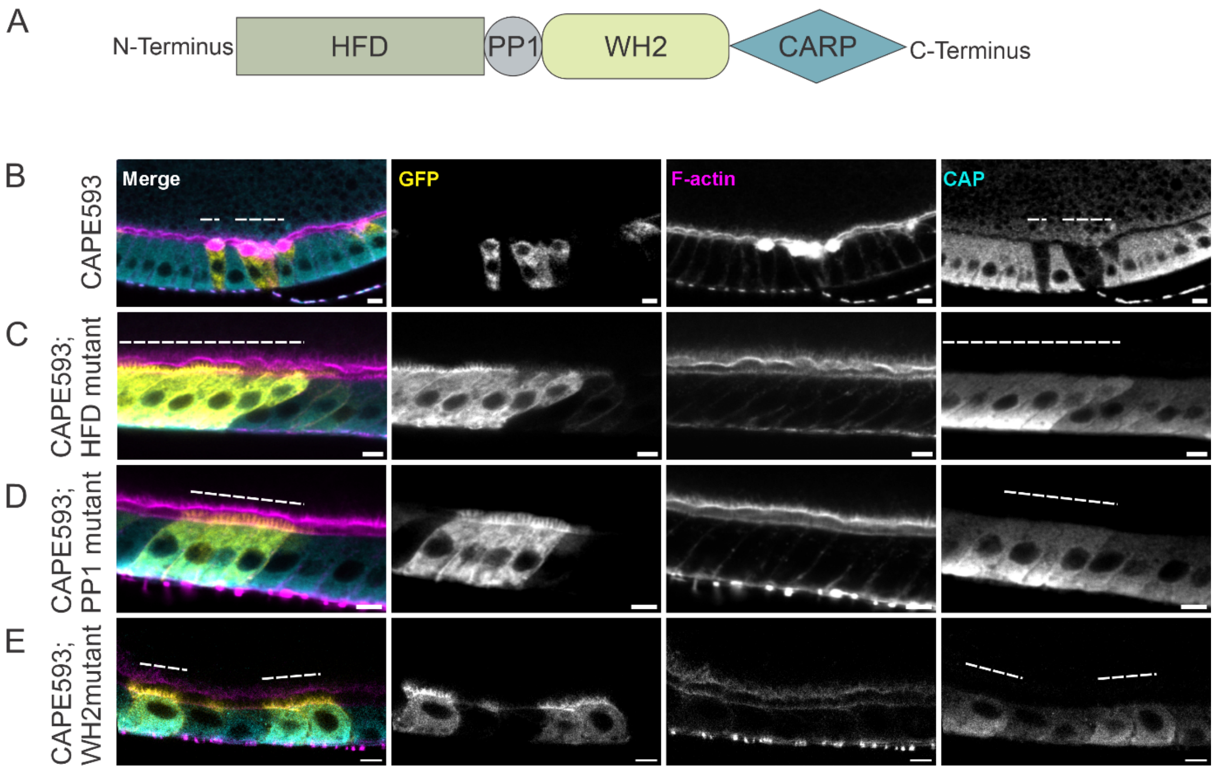


**Supplementary Figure 2. Rescue of apical actin turnover in CAP mutant cells. (A)** Schematic diagram of protein domain organisation of CAP. **(B)** Mosaic follicular epithelium with CAP mutant cells (GFP positive, pseudo-colored yellow, dashed line) exhibiting significant apical actin accumulation (Phalloidin staining, pseudo-colored magenta) alongside a loss of CAP expression (immunostaining, pseudo-colored cyan). Scale bar = 10 µm. **(C)** Mosaic follicular epithelium with CAP mutant cells (GFP positive, pseudo-colored yellow, dashed line) expressing CAP with a mutation in the HFD domain (immunostaining, pseudo-colored cyan). HDF mutant rescues apical actin accumulation (Phalloidin staining, pseudo-colored magenta) in CAP mutant cells. Scale bar = 10 µm. **(D)** Mosaic follicular epithelium with CAP mutant cells (GFP positive, pseudo-colored yellow, dashed line) expressing CAP with a mutation in the PP1 region (immunostaining, pseudo-colored cyan). PP1 mutant rescues apical actin accumulation (Phalloidin staining, pseudo-colored magenta) in CAP mutant cells. Scale bar = 10 µm.

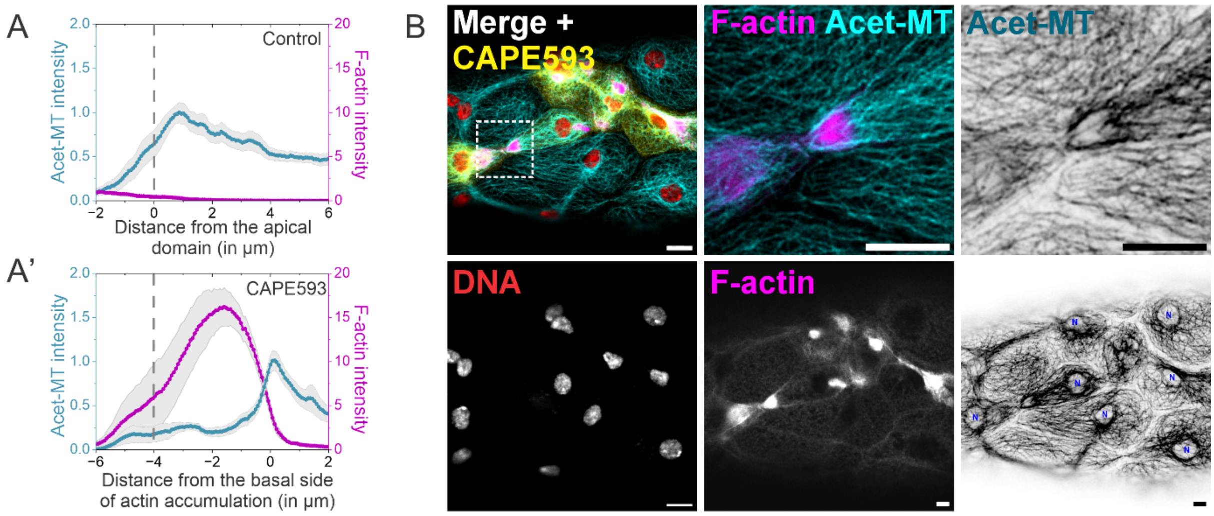


**Supplementary Figure 3. Actin accumulation in CAP mutant cells excludes microtubule. (A) Mean line scan intensity profile** of F-actin (magenta) and acetylated microtubules (cyan) is presented for control (A) and CAP mutant follicle cells (A'). In (A'), the x-axis represents the distance from the basal side of the accumulated F-actin to normalise for the variation in the space occupied by the accumulated actin in the cells. The dotted line marks the apical boundary of the follicle cells. Data are presented as mean ± SEM; n = 5 cells, N = 2 egg chambers. **(B) Actin accumulation excludes acetylated microtubules in CAP mutant stretch cells.** Stretch cells containing CAP mutant cells are marked by mCD8-GFP (GFP positive, pseudo-colored yellow). F-actin (phalloidin-stained, pseudo-colored magenta) accumulates at certain regions of the stretch cell cytoplasm. Zoomed-in inset (dotted box) highlights a site of filamentous actin accumulation (detected with Phalloidin, magenta) in the stretch cell is devoid of stable acetylated microtubules (Acet-MT, cyan). Additionally, acetylated microtubules are absent from the vicinity of the stretch cell nuclei, which are stained with DAPI (pseudo-colored red in the merged image and labelled "N" in the inverted greyscale LUT image). Scale bar = 10 µm.

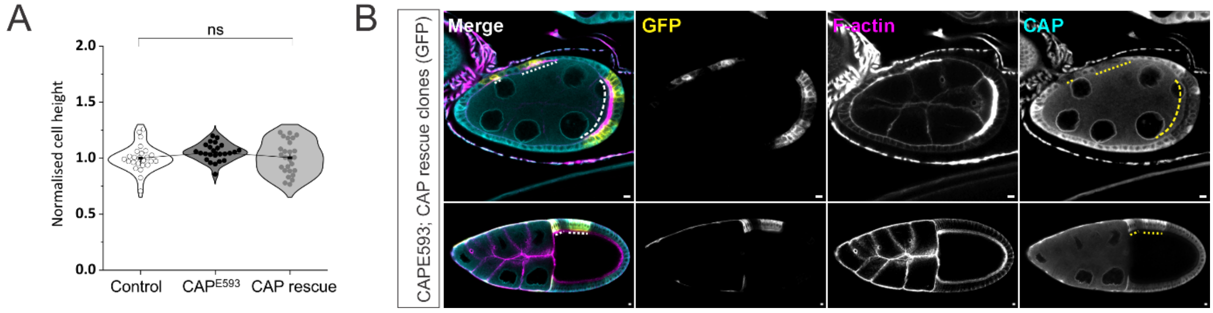


**Supplementary Figure 4: (A) Loss of CAP does not lead to change in cell height.** Quantification of cell height normalized with the control. The connecting line passes through the mean value of each group, and the whiskers represent the standard deviation. Kruskal–Wallis ANOVA followed by Dunn's post hoc test was used to assess the statistical significance of the different groups. Control = 25 cells, CAPE593 = 25 cells; CAP rescue = 26 cells. N = 5 egg chambers. **(B) Lack of CAP rescue construct expression during early oogenesis**.

Cross-sections of stage 7 (s7) and stage 10 (s10) egg chambers, with CAP rescue clones highlighted by mCD8-GFP (GFP positive, dashed line). The CAP rescue transgene is expressed (as indicated by immunostaining, pseudo-colored cyan) and rescues actin accumulation in stage 10. In stage 7, the CAP rescue construct is not expressed and thus fails to rescue the actin accumulation phenotype. Scale bar = 10 μm.

**Supplementary table 1: *Drosophila* genotypes used in supplementary figures**

| Figure | Panel | Genotype | Abbreviation | Source |
|---|---|---|---|---|
| S1 | A, C-E, G-H' | yw,hsFLP/w;TubGal80,FRT40A/Capt$^{E593}$ FRT40A; TubP-GAL4, UAS-mCD8-GFP / + | CAP$^{E593}$ | BDSC 5944 |
| | B | hsflp; UbimFRT40/Capt$^{E593}$FRT40A; AIP1GFP/+ | CAP$^{E593}$; AIP1GFP | BDSC 50824 |
| | F | hsFLP/y[1]P{w[+mC]=PTTGC}Actn[CC01961] w[*];UbimRFPnls,FRT40/Capt$^{E593}$ FRT40A | CAP$^{E593}$; α-Actinin | BDSC 602268 |
| | B | yw,hsFLP/w;TubGal80,FRT40A/Capt$^{E593}$ FRT40A; TubP-GAL4, UAS-Capt/ UAS-mCD8-GFP | CAP$^{E593}$; CAP Rescue | BDSC 5944 UAS-Capt, in this ms. |
| S2 | B | yw,hsFLP/w;TubGal80,FRT40A/Capt$^{E593}$ FRT40A; TubP-GAL4, UAS-mCD8-GFP / + | CAP$^{E593}$ | BDSC 5944 |
| | C | yw,hsFLP/w;TubGal80,FRT40A/Capt$^{E593}$ FRT40A; TubP-GAL4, UAS-Capt HFD/ UAS-mCD8-GFP | CAP$^{E593}$; HFD mutant | UAS-Capt, in this ms |
| | D | yw,hsFLP/w;TubGal80,FRT40A/Capt$^{E593}$ FRT40A;TubP-GAL4,UAS-CaptPP/ UAS-mCD8-GFP | CAP$^{E593}$; PP mutant | UAS-capt PP, in this ms |
| | E | yw,hsFLP/w;TubGal80,FRT40A/Capt$^{E593}$ FRT40A; TubP-GAL4, UAS-Capt mWH2/ UAS-mCD8-GFP | CAP$^{E593}$; WH2mutant | UAS-Capt mWH2, in this ms |
| S3 | B | yw,hsFLP/w;TubGal80,FRT40A/Capt$^{E593}$ FRT40A; TubP-GAL4, UAS-mCD8-GFP / + | CAP$^{E593}$ | BDSC 5944 |
| S4 | B | yw,hsFLP/w;TubGal80,FRT40A/Capt$^{E593}$ FRT40A; TubP-GAL4, UAS-Capt/ UAS-mCD8-GFP | CAP$^{E593}$; CAP Rescue | BDSC 5944 UAS-Capt, in this ms. |
